# MicroRNA-223 Enhances Microglia-Dependent Clearance of Amyloid Beta Plaques and Ameliorates Behavioral Deficits in a Mouse Model of Alzheimer’s Disease

**DOI:** 10.64898/2026.07.20.738977

**Authors:** Andre Krunic, Nikkitha Umesh Ganesh, Ugur Coskun, William Brennan, Chandani Patel, Ojasi Joshi, Jane Lee, Tori Shiyang Gu, Jaclyn Caruso, Aoife O’Connell, Charles Lisboa, Nicholas Crossland, Margareta Kurkela, TCW Julia, Abigail Fowler, Tuan Leng Tay, Andre Fischer, Ivana Delalle, Jan K. Blusztajn, Tiffany J. Mellott

**Affiliations:** Department of Pathology and Laboratory Medicine, Boston University Chobanian and Avedisian School of Medicine, Boston, MA, USA; Department of Pharmacology, Physiology, and Biophysics, Boston University Chobanian and Avedisian School of Medicine, Boston, MA, USA; Department of Epigenetics and Systems Medicine in Neurodegenerative Diseases, German Center for Neurodegenerative Diseases (DZNE), Göttingen, Germany; Department of Anatomy and Neurobiology, Boston University Chobanian and Avedisian School of Medicine, Boston, MA, USA; National Emerging Infectious Disease Lab, Boston University Chobanian and Avedisian School of Medicine, Boston, MA, USA; Bioinformatics Program, Faculty of Computing & Data Sciences, Boston University, Boston, MA, USA; Graduate Program in Genetics & Genomics, Boston University, Boston, MA, USA; Department of Biology, Boston University, Boston, MA, USA; Center for Systems Neuroscience, Boston University, Boston, MA, USA

## Abstract

The Alzheimer’s disease (AD) brain is characterized by dysregulated expression of multiple microRNAs (miRNA), positioning them as promising diagnostic and therapeutic targets. The levels of glia-enriched miR-223 are abnormal in the brains and plasma of AD patients and miR-223 is neuroprotective in models of stroke. However, whether miR-223 can be beneficial in AD is not known. Here, we report that intracerebroventricular (ICV) injection of miR-223 oligonucleotide mimic alleviated cognitive impairment, reduced amyloid beta (Aβ) pathology, and ameliorated the defects in synaptic marker expression in App^NL-G-F^ AD model mice. Mechanistically, miR-223 induced microglial clustering around Aβ plaques with a concomitant upregulation of microglial phagocytic receptors AXL, TREM2 and CD11c, while pharmacological microglial depletion abolished the plaque-clearance phenotype. Moreover, in human iPSC-derived microglia miR-223 directly targeted multiple genes in the endo-lysosomal pathway, including AD risk gene *SPPL2A*, indicating that it acts as a major regulator of microglial phenotype. Lastly, long-term AAV-mediated overexpression of miR-223 recapitulates its beneficial effects on cognition, pathology, and synaptic marker expression. Our study demonstrates a novel approach for the treatment of AD using miR-223 and highlights the potential of RNAi-based therapeutics in neurodegenerative disease.

## Introduction

Alzheimer’s Disease (AD) is the most common neurodegenerative disease worldwide. The pathological hallmarks of AD, amyloid beta (Aβ) plaques and tau tangles, synergistically drive downstream neuroinflammation, synaptic failure, neuronal loss, and eventual cognitive decline. MicroRNAs (miRNA) are short, 18-22 nucleotide (nt) long non-coding RNAs that negatively regulate gene expression through inhibition of their target mRNAs. The mature miRNA associates with Argonaute family proteins to form the RNA-induced silencing complex (RISC), which uses the miRNA to bind complementary mRNA sequences, typically in the 3’ untranslated region (UTR). In cases of full complementarity, the RISC cleaves its target mRNA, while in cases of partial complementarity it blocks translation, leading to de-adenylation and mRNA degradation^1^. Altered expression of miRNAs has been associated with all the molecular hallmarks of AD, including amyloidosis, tauopathy, inflammation, and synaptic dysfunction^2,3^. Furthermore, correcting the expression of dysregulated miRNAs has been shown to ameliorate AD pathophysiology in animal models. For instance, overexpression of the neuronal miR-132 leads to improved neurogenesis, while deletion of the microglial miR-155 enhances protective microglia polarization, demonstrating the potential of miRNAs as therapeutic targets in AD^4,5^.

miR-223 is a glia-enriched miRNA reported as neuroprotective in models of stroke and experimental autoimmune encephalomyelitis (EAE)^6–8^. Previous studies have shown that its levels are abnormal in plasma, CSF, and brain tissue of AD patients^9–18^. In a study of the Alzheimer’s Disease Neuroimaging Initiative (ADNI) cohort, CSF miR-223 levels were negatively associated with Aβ load in AD patients and its plasma levels were negatively associated with APOE4 status in the control subjects^19^ and our studies in the same cohort found that plasma levels of miR-223 were associated with an increased risk of progressing from mild cognitive impairment (MCI) to dementia^15^. Previous studies have shown that miR-223 promotes anti-inflammatory polarization of microglia in the rat brain^20^ and in mouse models of spinal cord injury and demyelination, miR-223 improved clearance of neuronal debris, while also reducing the burden of lipid droplets^21,22^. While miR-223 primarily regulates gene expression in glia, it is also secreted by these cells in exosomes, whose uptake into neighboring neurons results in modulation of their function, such as reduced hyperexcitability by targeting glutamate receptors and preventing neuronal apoptosis^6,23–25^.

Taken together, the evidence that AD pathophysiology is associated with reduced miR-223 expression and that miR-223 has therapeutic actions in models of neuronal injury provided a rationale for this study that aimed to assess the neuroprotective role of miR-223 in AD. Using the App^NL-G-F^ mouse model of amyloidosis, we found that miR-223 alleviated cognitive impairment, Aβ pathology, and ameliorated the defects in synaptic marker expression. The clearance of Aβ plaques promoted by miR-223 was abolished by microglia depletion indicating that it was dependent on these cells. In human iPSC-derived microglia, miR-223 positively regulated phagocytosis with transcriptomic evidence implicating the endo-lysosomal network in mediating its anti-amyloidogenic and anti-inflammatory actions. Lastly, long-term AAV-mediated overexpression of miR-223 demonstrated many of the same benefits as oligonucleotide mediated overexpression, including improved cognition, Aβ plaque clearance, reduced gliosis, and improved levels of synaptic markers.

## Results

### Restoring miR-223 Levels Reverses Cognitive Deficits in App^NL-G-F^ mice

miR-223 is found as two isotypes, miR-223-5p (passenger strand) and miR-223-3p (guide strand). miR-223-3p is the active form of the miRNA, and all reference to miR-223 herein will specifically refer to miR-223-3p. We first measured miR-223 levels in brain of wild type (WT) and App^NL-G-F^ model mice. In WT mice, miR-223 levels were approximately 3-fold higher in the hippocampus relative to the cerebral cortex (Figure 1A). In 7-month-old App^NL-G-F^ mice miR-223 levels were markedly decreased by 44% in the cortex and 24% in the hippocampus relative to WT controls at 7 months (Figure 1B), with a 63% decrease in the cortex and 51% decrease in the hippocampus at 12 months of age (Figure S1A-B).

**Figure 1:**
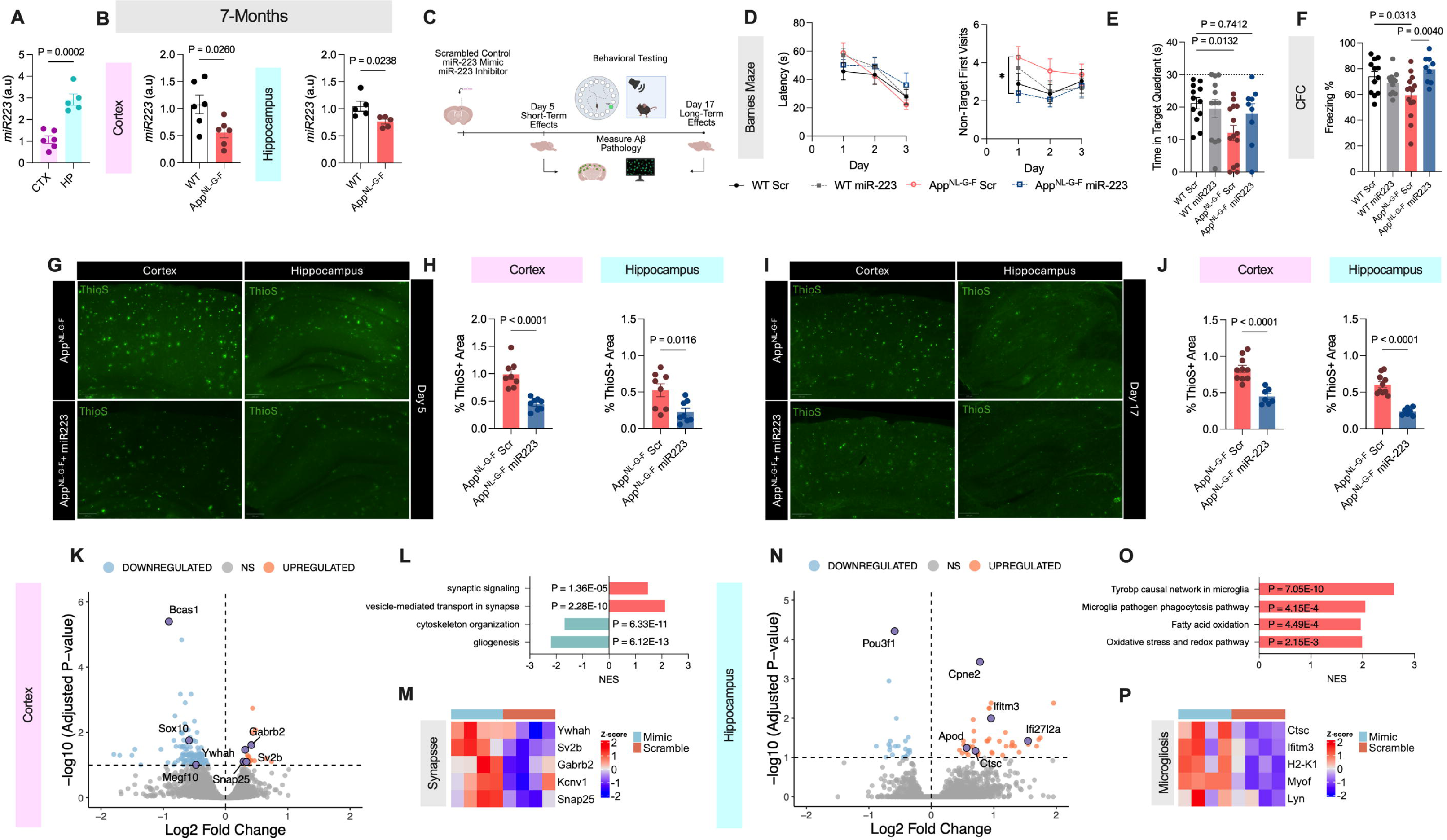
miR-223 Alleviates Behavioral Impairment and Amyloidosis in App^NL-G-F^ Mice. **A** Relative level of miR-223 in the cortex and hippocampus of wild type mice demonstrates biased expression in the hippocampus (n = 5-6 mice). **B** Levels of miR-223 between wild type and App^NL-G-F^ mice shows decreased expression in disease mice in the cortex (n = 6) and hippocampus (n = 5). **C** Experimental design of mouse studies. Mice were injected with either scrambled control, miR-223 mimic, or miR-223 inhibitor and were either sacrificed 5 days post injection or underwent behavioral testing and were sacrificed 17 days post injection. **D** Latency to target and non-target visits during Barnes Maze learning phase shows increased number of mistakes in AD mice that is reversed by miR-223 mimic administration (WT Scr: n = 12, WT miR-223: n = 12, App^NL-G-F^ Scr n = 14, App^NL-G-F^ miR-223 n = 9). **E** Time spent in target quadrant during the 24-hour probe test shows improved memory by miR-223. **F** Contextual fear conditioning (CFC) tone test freezing was impaired in App^NL-G-F^ mice and subsequently restored by miR-223. **G-H** Representative Thioflavin S (ThioS) images (**G**) and quantification (n = 8 mice) in the cortex and hippocampus **(H)** demonstrates robust reduction of amyloid beta pathology by miR-223 at 5 days post treatment. **I-J** Representative Thioflavin S (ThioS) images **(I)** and quantification (n = 7-10 mice) in the hippocampus and hippocampus **(J)** demonstrates robust reduction of amyloid beta pathology by miR-223 at 17 days post treatment. **K** Volcano plot of DEGs between miR-223 versus scrambled treated App^NL-G-F^ mice at 5 days post-treatment in the cortex. **L** Pathway enrichment in the cortex of day 5 miR-223 treated mice reveals upregulation for synaptic signaling. **M** Heatmap of select synaptic signaling genes in the cortex of day 5 miR-223 treated mice. **N** Volcano plot of DEGs between miR-223 versus scrambled control treated App^NL-G-F^ mice at 5 days post-treatment in the hippocampus. **O** Pathway enrichment in the hippocampus of day 5 miR-223 treated mice reveals upregulation for microglial phagocytosis and signaling. **P**Heatmap of select microglial genes in the hippocampus of day 5 miR-223 treated mice. Data shown as mean +/− SEM. **A-B, H, J** Two-tailed t-test, **D** three-way repeated measures ANOVA, **E** two-way ANOVA with Dunnett’s post-hoc test, **F** two-way ANOVA with Tukey’s post-hoc test.

Due to the reported neuroprotective actions of miR-223, we investigated whether altering its levels *in vivo* could affect AD-associated pathology. Seven-month-old female and male WT or App^NL-G-F^ mice were injected ICV with lipid nanoparticles containing 0.25 nmol/µL of miR-223 mimic, inhibitor (an antisense oligonucleotide designed to deplete the endogenous pool of miR-223), or scrambled control, followed by behavioral testing (Figure 1C). Seven months was chosen as this time point is when App^NL-G-F^ mice begin to show hippocampal pathology and behavioral deficits, with the early time point making the mice most amenable to treatment^26–28^. At 5 days post injection, miR-223 levels were increased by 3-fold in the cortex, demonstrating its successful overexpression (Figure S1C). Additionally, we found a trend towards increased miR-223 levels at the end of behavioral testing at post-injection day 17, demonstrating a relatively long half-life of the injected oligonucleotide (Figure S1C). In the open field test, we found no differences in average velocity due to oligonucleotide injection relative to a no-surgery sham control, demonstrating that mice recovered from the injection with no overt issues (Figure S1D). Moreover, we found that App^NL-G-F^ mice treated with a miR-223 inhibitor demonstrated a trend towards less exploratory behavior compared to scrambled treated App^NL-G-F^ mice, suggesting increased anxiety in the former (Figure S1D). This was not observed in WT mice. Next, we assessed spatial learning and memory using the Barnes Maze. All mice successfully learned the location of the escape hole, with App^NL-G-F^ mice not yet showing impairment in latency to hole presumably due to the relatively early stage of hippocampal pathology at 7 months of age (Figure 1D, S1E-H). However, App^NL-G-F^ mice made a greater number of non-target visits relative to WT mice, which was prevented by miR-223 mimic administration (Figure 1D). Likewise, treatment with miR-223 inhibitor did not change the number of errors made by App^NL-G-F^ mice (Figure S1F). Moreover, miR-223 improved performance on the 24-hour probe test in App^NL-G-F^ mice (Figure 1E). Unexpectedly, we found that miR-223 inhibitor treatment also partially improved latency and time in target zone on the probe test (Figure S1I-K). Because Barnes maze escape is motivated by aversion to the exposed bright and noisy environment, enhanced anxiety observed in the open field in the inhibitor-treated mice may be driving this effect.

We further probed fearful learning and memory using the contextual fear conditioning (CFC) test. Following the learning phase (day 1), mice were tested for freezing in response to the tone (day 2) or shock context (day 3). All mice were able to associate the tone with the shock, with miR-223 mimic-treated App^NL-G-F^ mice showing a trend towards faster learning (Figure S1L). On the tone test, App^NL-G-F^ mice were impaired and miR-223 mimic treatment reversed this deficit in fearful learning (Figure 1E). Inhibitor treatment did not influence the tone response (Figure S1M). On the context test, miR-223 mimic or inhibitor treatment did not influence freezing response (Figure S1N).

### miR-223 Ameliorates Amyloidosis in App^NL-G-F^ mice

Based on our behavioral phenotype, we next assessed whether miR-223 could reduce amyloid burden in brain of App^NL-G-F^ mice. At 5 days post-injection, there was a 56% decrease in Thioflavin S (ThioS) positive amyloid staining area in the cortex and a 66% decrease in the hippocampus (Figure 1G-H). Extending to 17 days post-injection, this reduction in ThioS+ area persisted in both brain regions, with a 47% decrease in the cortex and 72% decrease in the hippocampus (Figure 1I-J), demonstrating that miR-223 robustly and rapidly reduces amyloid burden in App^NL-G-F^ mice. To validate our findings with an alternative approach, we stained for dense-cored Aβ plaques using AmyloGlo and quantified plaque density using ImageJ. In the cortex, miR-223 reduced the number of plaques per unit area (Figure S2A-C). Additionally, the hippocampus showed a trend towards fewer plaques measured by AmyloGlo. Treatment with miR-223 inhibitor did not alter amyloid pathology in either region when measured by AmyloGlo or ThioS (Figure S2A-E).

### miR-223 Engages Microglial and Neuronal Signaling Pathways *In vivo* and Attenuates AD-associated Changes

To gain mechanistic insights into the functions of miR-223 *in vivo*, we performed bulk RNA sequencing on the cortex and hippocampus of mice treated with miR-223 or scrambled control oligonucleotide at 5 days and 17 days post injection. Target engagement was assessed using GSEA with 308 experimentally validated miR-223-3p targets based on the miRTarBase database^29^. In the cortex, there was a trend towards decreased expression of miR-223 targets, however this was not statistically significant (Figure S3A). In the hippocampus, we found a substantial downregulation of miR-223 target expression, confirming direct target engagement following mimic treatment (Figure S3A). At day 17, we found that miR-223 validated target genes were significantly downregulated in the cortex (Figure S3B). However, we found a strong induction of miR-223 targets in the hippocampus (Figure S3B), possibly due to rebound effects that have been reported when using RNA interference (RNAi) *in vivo*^30,31^.

We next characterized gene and pathway-level changes at individual time points. At day 5 in the cortex, we found 143 DEGs when comparing miR-223 to scrambled control (Figure 1K, Figure S3C). On the pathway level, there was a significant upregulation of synaptic vesicle transport and synaptic signaling, with a downregulation in cytoskeleton and gliogenesis genes (Figure 1L). Notable genes involved in synaptic signaling induced by miR-223 include *Snap25, Ywhah, Gabrb2, Kcnv1, and Sv2b*, demonstrating an overall pro-synaptic phenotype (Figure 1M). At day 17 in the cortex, there were no genes meeting our FDR cutoff (FDR < 0.1), possibly due to the overall small effect size on individual gene expression elicited by miR-223 as well as the relatively long period post-treatment (Figure S3C, E). However, miR-223 treatment reduced the expression of genes in pathways related to cytokine production and the adaptive immune response (Figure S3F). To capture more subtle gene-level changes, we assessed genes that were significantly altered between WT:Scr and App^NL-G-F^:Scr mice (FDR < 0.1) that are not significant when comparing WT:Scr and App^NL-G-F^:miR-223. Using this method, we found that miR-223 treatment has an overall dampening effect on genotype-related DEGs (WT:Scr and App^NL-G-F^:Scr DEGs = 682, WT:Scr and App^NL-G-F^:miR223 = 125, Figure S3D), which we annotate as “miR-223-protected DEGs”. Notable upregulated genes that were partially normalized by mimic treatment included *Aif1* (encoding IBA1), lysosomal protein *Idua*, and tyrosine kinase *Fes*. Downregulated genes that were restored by miR-223 treatment included inhibitor of inflammatory signaling *Nfkbia*, as well as the Adiponectin receptor *Adipor2* (Figure S3G).

At day 5 in the hippocampus, there were 59 DEGs when comparing miR-223 to scrambled control (Figure 1N) with an overall increase in microglial phagocytosis and Tyrobp signaling (Figure 1O) in the mimic group relative to control. Individual genes within this pathway included tyrosine kinase *Lyn*, lysosomal component *Ctsc*, interferon response gene *Ifitm3,* and MHCII gene *H2-K1* (Figure 1P). The most upregulated gene in the hippocampus of miR-223 treated mice was *Myof*, a membrane fusion gene possibly involved in membrane repair^32^. At day 17 in the hippocampus, we surprisingly found the largest number of DEGs with 150 genes changed between miR-223 and scrambled control App^NL-G-F^ mice (Figure S3C-D, H). Notably, upregulated genes were strongly associated with neuron projection development, while we observed a downregulation in aerobic respiration (Figure S3I). As in the cortex, miR-223 had an overall dampening effect on genotype-related DEGs (WT:Scr and App^NL-G-F^:Scr DEGs = 2812, WT:Scr and App^NL-G-F^:miR223 = 1140, Figure S3C, D). Notable genes related to neuronal function upregulated by miR-223 include *Lrp8,* encoding the APOER2 receptor for Reelin, *Insr, Shisa6, Bmpr2,* and *Celsr3* (Figure S3J).

Lastly, to compare overall transcriptome-wide effects of miR-223, we correlated gene expression changes at day 17 between the effect of genotype (WT:Scr vs App^NL-G-F^:Scr) and miR-223 treatment (App^NL-G-F^:Scr vs App^NL-G-F^:miR-223). In both the cortex and hippocampus, we found that changes in gene expression due to miR-223 were negatively correlated with those due to genotype (Figure S3K), demonstrating an overall reversal of the disease-associated transcriptome by miR-223.

### miR-223 Restores Synaptic Markers in App^NL-G-F^ Mice

As miR-223 ameliorated behavioral deficits, reduced amyloid pathology and induced the expression of multiple neuronal genes, we next assessed whether this was associated with normalized expression of synaptic marker proteins by performing 6-plex IHC with antibodies against Aβ42, dystrophic neurite marker LAMP1, pre- and postsynaptic markers SYP and SAP102, cholinergic marker ChAT, and neuronal marker MAP2 (Figure S4A). In App^NL-G-F^ mice, we found an increase in LAMP1+ dystrophic neurite area, as well as reduced area of SYP, ChAT, and MAP2 in the cortex (Figure 2A). Using antibody staining, we validated the histochemical data that miR-223 reduced Aβ area in the cortex (Figure 2B). Parallel to the decrease in Aβ, we found that miR-223 significantly reduced LAMP1+ area in the cortex, indicating reduced neuronal damage (Figure 2C). In addition, miR-223 increased SYP levels in App^NL-G-F^ mice (Figure 2D), while there was a trend towards increased ChAT (Figure 2E) and MAP2 (Figure 2F) area staining. SAP102 levels were not significantly changed by genotype, however, were elevated relative to WT control in the miR treated group, suggesting that miR-223 can alter post-synaptic protein levels independently of reversing AD-associated changes (Figure 2G).

**Figure 2:**
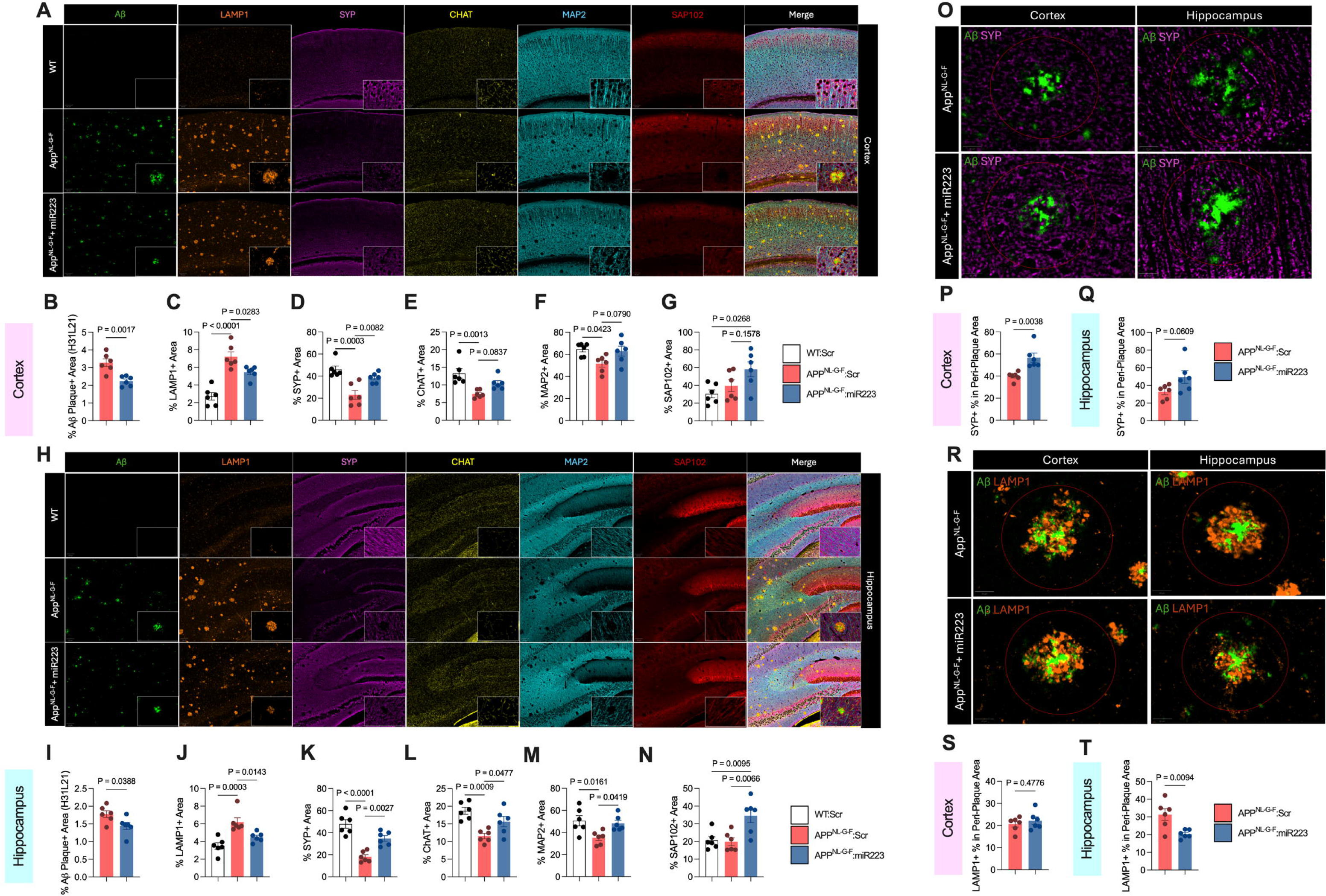
miR-223 Increases Levels of Synaptic Markers and Reduces Dystrophic Neurites. **A** Representative multiplex immunohistochemistry (IHC) images of the cortex stained for Aβ, LAMP1, SYP, CHAT, MAP2, and SAP102. **B-G** Percent area quantification (n = 6) in the cortex of Aβ **(B)**, LAMP1 **(C)**, SYP **(D)**, CHAT **(E),** MAP2 **(F)**, and SAP102 **(G)** demonstrates reduction in LAMP1+ dystrophic neurites and Aβ with subsequent restoration of synaptic markers. **H** Representative multiplex immunohistochemistry (IHC) images of the hippocampus stained for Aβ, LAMP1, SYP, CHAT, MAP2, and SAP102. **I-N** Percent area quantification (n = 6) in the hippocampus of Aβ **(I)**, LAMP1 **(J)**, SYP **(K)**, CHAT **(L),** MAP2 **(M)**, and SAP102 **(N)** demonstrates reduction in LAMP1+ dystrophic neurites and Aβ with subsequent restoration of synaptic markers. **O-Q** Representative images **(O)** and quantification in the cortex **(P)** and hippocampus **(Q)** of peri-plaque SYP staining shows increased synaptic marker levels in the cortex despite presence of Aβ pathology. Each data point represents the average of 5 plaques per animal**. R-T** Representative images **(R)** and quantification in the cortex **(S)** and hippocampus **(T)** of peri-plaque LAMP1 staining shows decreased dystrophic neurite severity in the hippocampus despite presence of Aβ pathology. Each data point represents the average of 5 plaques per animal. Data shown as mean +/− SEM. **B-G, I-N** one-way ANOVA with Tukey’s post-hoc test. **P,Q,S,T** two-tailed t-test.

In the hippocampus, as in the cortex, we observed an increase in LAMP1+ dystrophic neurites in App^NL-G-F^ mice, with subsequent decreases in SYP, ChAT, and MAP2 positive areas (Figure 2H). When measured by IHC, miR-223 decreased Aβ staining area (Figure 2I), while concurrently decreasing LAMP1+ area (Figure 2J). Moreover, miR-223 increased levels of SYP (Figure 2K), ChAT (Figure 2l) and MAP2 (Figure 2M) positive areas. Like in the cortex, SAP102 levels did not change between WT and App^NL-G-F^ mice, however miR-223 increased SAP102 staining areas beyond WT levels (Figure 2N).

Synapses are locally depleted in the vicinity of Aβ plaques^33^, with presynaptic sites showing specific vulnerability^34^. Therefore, we quantified SYP staining within the immediate area of Aβ plaques (Figure 2O). In the cortex, there was an increase in SYP+ area in miR-223 treated App^NL-G-F^ mice as compared to scrambled controls in the plaque vicinity and a trend towards increasing in the hippocampus when averaging per animal (Figure 2P-Q), demonstrating increased synaptic marker levels despite local presence of Aβ pathology. When comparing the distribution of per-plaque SYP+ area, we found an overall increase in both the cortex and hippocampus (Figure S4B-C). We also quantified LAMP1 area around plaques to determine whether miR-223 can reduce dystrophic neurites in the presence of Aβ (Figure 2R). There was no change in LAMP1+ staining in the area around Aβ plaques in the cortex, suggesting that the overall reduced LAMP1+ area in the cortex is due to a reduced number of plaques (Figure 2S, Figure S4D). In contrast, LAMP1+ staining was reduced in the peri-plaque area in the hippocampus, demonstrating that miR-223 reduced neuronal injury despite the presence of Aβ (Figure 2T, Figure S4E).

### miR-223 Promotes Microglial Engagement of Amyloid and Increases Levels of Phagocytic Proteins

In response to Aβ, microglia exhibit an inflammatory response and cluster around plaques to create a physical barrier between Aβ and the surrounding parenchyma. This is coupled with a change in morphology as microglia switch from a ramified to amoeboid state and contribute to Aβ clearance via phagocytosis. Based on the observed reduction of amyloidosis in miR-223-treated App^NL-G-F^ mice and the predominantly glial expression of this miR, we hypothesized that miR-223 can alter the response of microglia to Aβ. To test this hypothesis, we used AutoMorFi to quantify microglial morphology on IBA1-stained brain sections after 5 days of miR-223 treatment^35^. We found an increase in microglial process length in the cortex suggesting an overall more ramified state (Figure 3A). This was not seen in the hippocampus, in line with the pro-phagocytic and amoeboid mRNA changes observed previously (Figure 3A, Figure 1O). To validate our findings, we performed Sholl analysis on microglia from both regions and found an increase in the number of intersections in the cortex of mimic treated mice (Figure S5A). We followed this by quantifying microglial morphology at day 17 post treatment (Figure S5B). We found that in App^NL-G-F^ mice, there was an overall loss of microglial ramification and increased clustering in the cortex in response to pathology, however we did not see a change in ramification with miR-223 treatment (Figure S5C-E).

**Figure 3:**
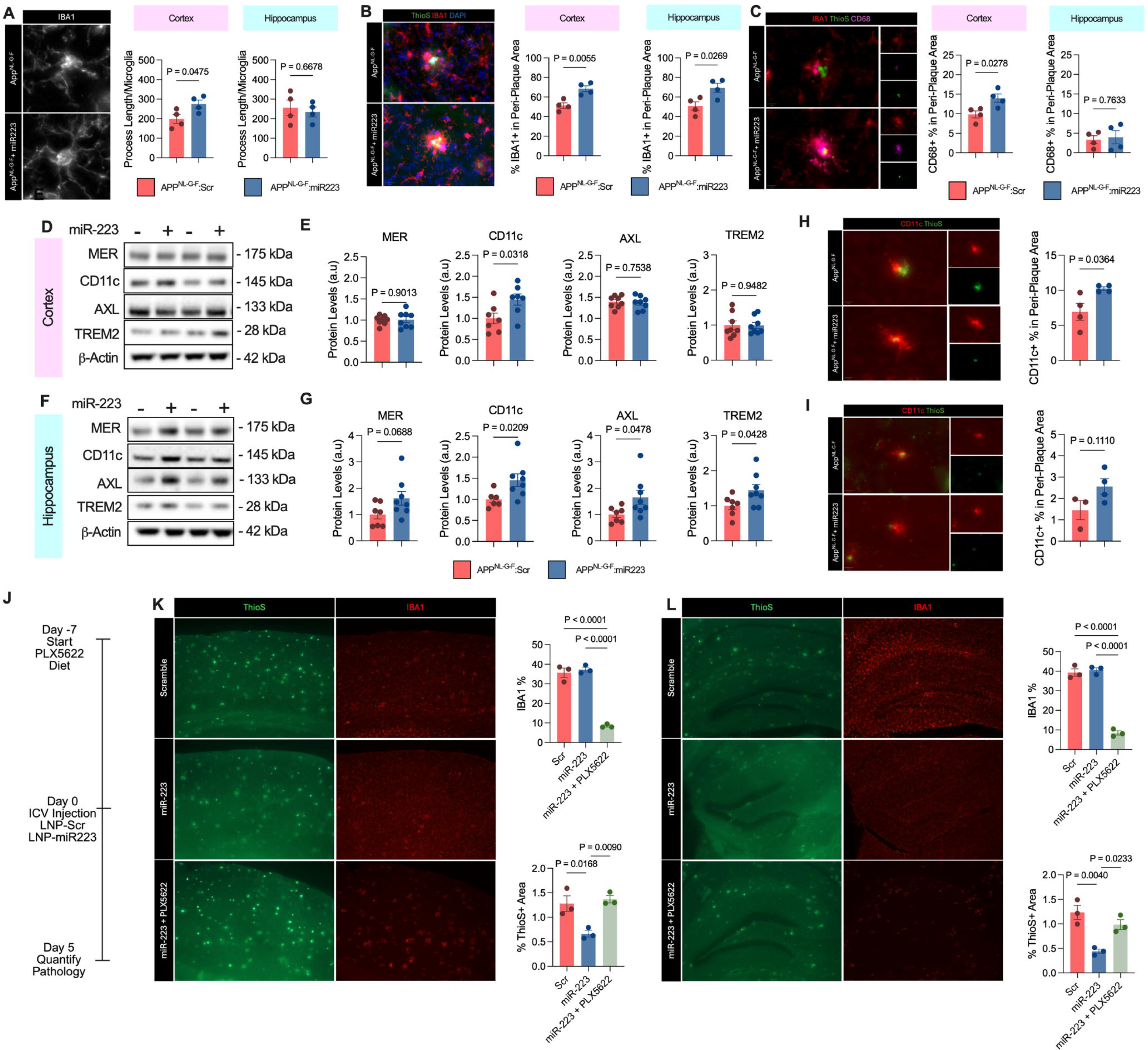
miR-223 Induces Microglial Engagement of Amyloid. **A** Representative microglia in the cortex of miR-223 treated mice and quantification of average process length in the cortex and hippocampus. **B** Representative image of IBA1+ area in the vicinity of amyloid beta plaques and quantification in the cortex and hippocampus shows enhanced microglial engagement of plaques in miR-223 treated mice. **C** IBA1/CD68 co-staining representative image and quantification in the cortex. **D-E** Representative western blotting and quantification of microglial receptors TREM2, MER, CD11c and AXL in the cortex of miR-223 treated mice. **F-G** Representative western blotting and quantification of microglial receptors TREM2, MER, CD11c and AXL in the hippocampus of miR-223 treated mice. **H-I** Immunofluorescence representative images and quantification of Aβ-associated CD11c+ area in the cortex and hippocampus. **J** Mice were fed a diet containing PLX5622 for 7 days prior to miR-223 or scrambled control treatment and ThioS area was assessed to determine plaque load. **L-M** Representative images of ThioS+ plaque area and IBA1 staining and quantification in the cortex of PLX5622 treated mice reveals ablation of plaque clearance by miR-223 in microglia-depleted mice. +/− SEM. **B-D, F, H, I-J** two-tailed unpaired t-test. **L, M** one-way ANOVA with Tukey’s post-hoc test.

Next, assessed how miR-223 altered the microglial marker response to Aβ. We found that miR-223 increased IBA1+ signal surrounding Aβ plaques in both the cortex and hippocampus at 5 days post treatment (Figure 3B). Moreover, this effect was still present at day 17 post treatment in the cortex, while diminishing in the hippocampus (Figure S5F). We than co-stained the tissue sections for IBA1 and CD68, a lysosomal protein associated with activated microglia. In the cortex, miR-223 increased CD68+ microglia area surrounding plaques, suggesting increased lysosomal processing (Figure 3C). However, we found no difference in microglial peri-plaque CD68 in the hippocampus (Figure 3C). It should be noted that CD68 immunoreactivity in the hippocampus was lower than in the cortex overall, likely due to the earlier stage of pathology. We next quantified the levels of several proteins associated with the microglial response to Aβ at 5 days post miR-223 treatment. These included levels of cell surface receptors involved in Aβ recognition and clearance, namely AXL, MER, CD11c, and TREM2 (Figure 3D). In the cortex, miR-223 increased CD11c protein levels by 45%, while we found no difference in TREM2, MER, and AXL (Figure 3E). Notably, CD11c is a microglial marker associated with protection against Aβ^36,37^. In the hippocampus, we observed a 44% increase in TREM2 and a 46% increase in CD11c due to miR-223 treatment (Figure 3F, G). We also found that levels of AXL were increased by 65% and while MER showed a trending increase of 62% (Figure 3F, G). Taken together, these data demonstrate that miR-223 promotes an amyloid pro-clearance phenotype in microglia, possibly by upregulating a subset of relevant receptors. To determine whether these changes persisted, we assessed protein levels at day 17. At this time point however, we found that only the levels of AXL in the cortex were significantly increased, with other receptors showing no change (Figure S5G). In addition to phagocytic receptors, we assessed levels of microglial activation markers 5days following miR-223 administration, namely the NLRP3 inflammasome. In the cortex, adapter protein ASC was significantly downregulated by treatment, however NLRP3 itself did not change (Figure S5H). In the hippocampus, the protein levels of NLRP3 inflammasome components did not change (Figure S5I). To validate the relative contribution of CD11c+ microglia towards Aβ clearance by miR-223, we performed immunofluorescence to determine the localization of CD11c+ cells in the brain. Immunostaining revealed that CD11c+ microglia were predominantly located around Aβ plaques, in line with its status as a disease-associated microglia (DAM) gene (Figure S5J). We found a significant increase in CD11c positive area in the Aβ plaque area in the cortex (Figure 3H) as well as a trending increase in the hippocampus (Figure 3I).

### Microglia are Required for miR-223-mediated Clearance of Amyloid

We next assessed the causal role of microglial clearance in mediating the anti-Aβ effect of miR-223. Mice were fed either a control or PLX5622-containing diet for 7 days prior to ICV injections with miR-223 mimics or scrambled controls (Figure 3J). PLX5622 is a specific CSF1R inhibitor that rapidly depletes microglia in the brain and indeed we found that PLX5622 treatment greatly reduced microglia number in the cortex and hippocampus (Figure 3K, L) and did not itself substantially influence plaque burden following our exposure period (Figure S5K). In line with the above-data (Figure 1G-J), treatment with miR-223 itself reduced plaque area assessed by ThioS in both the cortex and hippocampus; however, this effect was fully ablated by pre-exposure to PLX5622 (Figure 3K, L) indicating that miR-223 enhances the removal of Aβ plaques through microglial-dependent phagocytic clearance.

### miR-223 Regulates Phagocytosis and Inflammation in Human and Murine Microglia

To further establish the role of miR-223 in microglial function *in vitro*, we assessed phagocytic capacity of human iPSC-derived microglia (iMG) transfected with either a miR-223 mimic or inhibitor (Figure 4A). Overexpression of miR-223 enhanced (Figure 4B), while knock-down of miR-223 inhibited phagocytosis (Figure 4C), demonstrating that miR-223 levels can bi-directionally control phagocytic activity. To further validate the effects of miR-223 on phagocytosis, we transfected mouse primary microglia with miR-223 (Figure S6A-B), and confirmed the engagement of its direct targets, including decreased *Il6* and *Nlrp3* expression, as well as induction of *Itgax* (the gene encoding CD11c) (Figure S6C), the latter replicating our *in vivo* findings. In line with its effect on human microglia, we found that miR-223 increased phagocytosis in primary microglia (Figure S6D).

**Figure 4:**
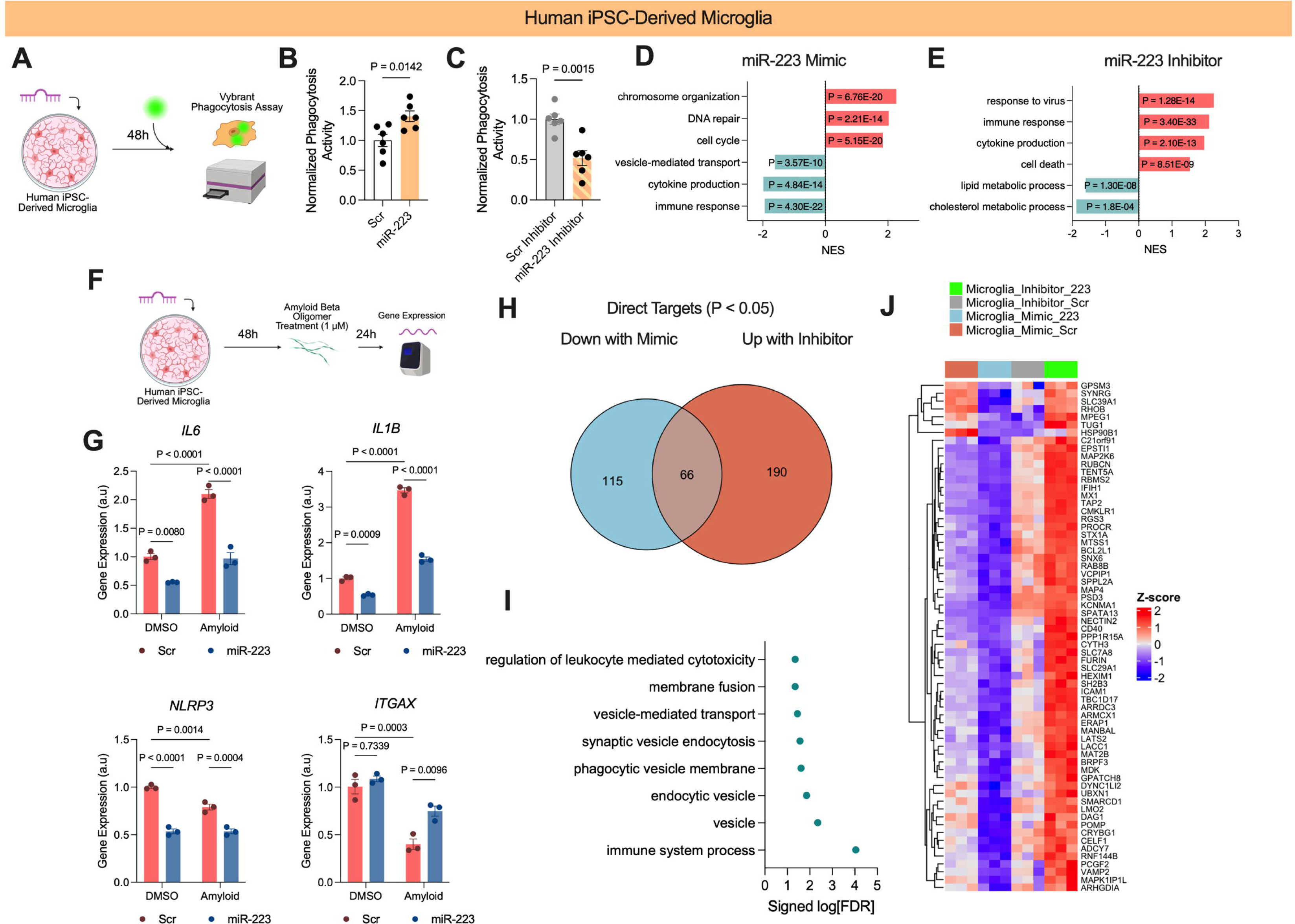
miR-223 is a Direct Regulator of Microglial Phagocytosis. **A** Experimental design for phagocytosis in iPSC-derived microglia (iMG). **B-C** Normalized phagocytic activity in iMGs treated with miR-223 mimic (**B**) or inhibitor (**C**). **D-E** Pathway enrichment of upregulated and downregulated DEGs changed by 48-hour miR-223 mimic (**D**) or inhibitor (**E**) in iMG cells. **F** Experimental design for miR-223 and Aβ oligomer (Aβo) co-treatment. **G** Gene expression for microglial genes *IL6, IL1B, NLRP3*, and *ITGAX* with miR-223 and Aβo. **H** Overlap of direct targets of miR-223 downregulated by miR-223 mimic treatment and de-repressed by miR-223 inhibitor treatment reveals common targets. **I-J** Pathway enrichment (**I**) and heatmap (**J**) of 66 common genes associated with mimic and inhibitor treatment in iMGs.

We next conducted RNA-seq on iMGs treated with either a miR-223 mimic or inhibitor to further characterize the molecular pathways engaged by miR-223. The overexpression of miR-223 was validated by qPCR (Figure S6E). We identified 864 downregulated and 730 upregulated DEGs (Padj < 0.05), with downregulated pathways strongly enriched for immune and inflammatory processes and upregulated pathways enriched for cell cycle-related genes (Figure 4D, Figure S6F). Specific genes associated with immunity downregulated by miR-223 include the human leukocyte antigen (HLA) family, *C1QA-C, IL1B,* and *CD74* (Figure S6G), the latter of which is upregulated in AD-associated microglia in humans^38^.

Conversely, treatment with a miR-223 inhibitor markedly reduced endogenous miR-223 levels in iPSC-derived microglia (Figure S6H). In turn, we observed an overall increase in immune signaling and cytokine mRNA levels with miR-223 inhibition, including increased expression of *HLA, C1QA-C*, *CSF1, CXCL10, and IFIT2* (Figure 4E, Figure S6I-J). Homeostatic microglial markers *CX3CR1* and *P2RY12*, as well as *ITGAX* were also downregulated by miR-223 inhibition, suggesting an overall shift towards microglia activation evoked by this treatment (Figure S6J).

### miR-223 Alleviates Amyloid Beta Oligomer-Induced Inflammation in Human iMGs

To determine if miR-223 could directly counteract Aβ-associated changes in human cells, we treated iMGs with miR-223 for 48h, followed by treatment with 1 µM of Aβ oligomers (Aβo) for 24h (Figure 4F). Aβo treatment robustly increased mRNA levels of cytokines *IL6* and *IL1B*, confirming an inflammatory response (Figure 4G). Treatment with a miR-223 mimic attenuated these responses to Aβo (Figure 4G). Likewise, we found that the expression of miR-223 direct target, NLRP3, was reduced in both the control-(DMSO) and Aβo-treated cells (Figure 4G). Lastly, we assessed mRNA levels of *ITGAX* to confirm if our *in vivo* phenotype could be replicated in human cells. At baseline, miR-223 did not elevate *ITGAX*, however, Aβo treatment significantly decreased *ITGAX* mRNA and overexpression of miR-223 attenuated this effect, suggesting that miR-223 increases *ITGAX* expression selectively in pro-inflammatory microglia (Figure 4G). Treatment with miR-223 inhibitor significantly increased *IL6* expression under baseline conditions, however, did not significantly change *IL6* mRNA levels in Aβo treated cells (Figure S6K). Surprisingly, miR-223 inhibitor also attenuated Aβo-induced expression of *IL1B* (Figure S6K). miR-223 inhibitor had no effect on *NLRP3*, expression, while *ITGAX* levels were decreased by miR-223 inhibitor in unstimulated microglia, but not affected by it in with Aβo-treated cells (Figure S6K).

### miR-223 Directly Targets Genes Involved in the Endo-Lysosomal pathway

We next investigated direct targets of miR-223 to identify potential mechanisms underlying the observed phenotypes in human iMGs. Due to the repressive role of miRNAs on gene expression, we focused specifically on direct target genes compiled from 4 miRNA target databases (miRTarBase, miRDB, miRWalk, and Targetscan)^29,39–41^ that were either downregulated with mimic treatment or upregulated with inhibitor treatment. Using these criteria, we found 181 direct targets that were downregulated with mimic treatment and 256 targets that were upregulated with inhibitor treatment, with 66 of these being overlapping (Figure 4H). Performing pathway analysis of overlapping targets, we found an enrichment for endosome and phagosome-associated genes, such as *MPEG1*, *MTSS1*, *RUBCN*, *SPPL2A*, and *CD40* (Figure 4I, J). Analysis of all direct targets downregulated by miR-223 mimic treatment identified additional endosomal genes such as *INPP5B* and *RAB10,* being downregulated in the mimic-treated group only (Figure S6L, M). Direct targets upregulated by inhibitor treatment were more strongly associated with apoptosis and cytokine signaling, such as *CCL3*, *IFIH1*, *MX1*, *STAT1*, and *IL6*, however they were also significantly enriched for endosomes, including unique genes such as *CLN3*, *SNX6*, and *SNX30* (Figure S6N, O).

### miR-223 Alters Neuronal Signaling in Human iPSC-Derived Neurons

In addition to our *in vivo* studies showing major effects of miR-223 administration on the function of microglia, the treatment also resulted in neuroprotection (Figures 1M, Figure 2) and amelioration of cognitive defects in the AD model mice (Figure 1D-F) indicating that miR-223 may be active in neurons. To test this idea, we transfected human iPSC-derived neurons (iNeurons) with either a miR-223 mimic or inhibitor and performed bulk RNA-sequencing. Overexpression of miR-223 was confirmed via qPCR (Figure S7A). We found that miR-223 robustly altered gene expression, with 1705 downregulated and 1204 upregulated DEGs (Padj < 0.05, Figure S7B). At the pathway level, upregulated genes were enriched for vesicle organization, as well as autophagy (Figure S7C). Likewise, cell death, and ATP synthesis, and chromosome organization were enriched in downregulated DEGs (Figure S7C). Upregulated synaptic genes included glutamate receptors *GRIN2B* and *GRIA1*, voltage gated calcium channel *CACNA1B*, presynaptic components *SYNJ1* and *SYN2* and post-synaptic components *SHISA9, EPHB2, and DLGAP2* (Figure S7D). Treatment with a miR-223 inhibitor caused an almost total knock-down of miR-223 in iNeurons (Figure S7E). Despite adequate knockdown, treatment with a miR-223 inhibitor elicited relatively fewer changes in gene expression, with 422 downregulated and 398 upregulated DEGs (Padj < 0.05, Figure S7F). Moreover, DEGs identified in inhibitor treated iNeurons were small in magnitude, suggesting miR-223 may be partially dispensable in neurons. Inhibitor treatment showed a general reverse trend to that of the mimic, with downregulation of autophagy and vesicle transport (Figure S7G). Additionally, mature neuronal markers genes such as *NEFL* and *MAPT* were downregulated at the mRNA level (Figure S7H).

We next assessed what specific direct targets were altered by miR-223 mimic or inhibitor treatment in iNeurons. Treatment with mimic repressed several targets, including *PARP1*, which was previously proposed as a therapeutic target for inhibition in AD (Figure S7I)^42^. One notable target upregulated by inhibitor treatment was *HDAC4*, with HDAC inhibitors generally demonstrating a neuroprotective function (Figure S7J)^43^. Pathway enrichment of direct targets altered in iNeurons revealed a general enrichment for neuron differentiation and cell division (Figure S7K-L). Interestingly, only 7 genes were reciprocally regulated by mimic and inhibitor treatment, with the relatively fewer targets observed with the inhibitor, consistent with its low endogenous neuronal expression and with the notion that miR-223 may be partially dispensable for neuronal gene regulation (Figure S7M, N). To assess how miR-223 alters neuronal functions, we performed calcium flux assay on iNeurons treated with miR-223 mimic or inhibitor. Overexpression of miR-223 increased calcium influx upon glutamate treatment, while miR-223 inhibition showed a trend towards decrease calcium flux (Figure S7O). Taken together our data show that miR-223 promotes neuronal identity in human iPSC-derived neurons and support the notion that some of its disease modifying beneficial effects in App^NL-G-F^ mice may include these neurotrophic actions.

### Long Term Viral-Mediated Overexpression of miR-223 Reduces Amyloidosis and Inflammation and Increases Levels of Synaptic Markers

The experiments presented above utilizing single ICV injections of the mimic in AD model mice indicate that miR-223 has potential as a therapeutic agent for AD. However, the single injection approach produces only transient effects, not suitable for a chronic condition like AD, where repeated dosing is not practical. For this reason, we delivered miR-223 via a viral vector that provides durable expression to better model long-term target engagement. Following ICV injection of an AAV containing the endogenous mouse miR-223 sequence (or scrambled control sequence) in 6-month-old App^NL-G-F^ mice, we assessed their behavior and pathology 3 months later, at 9-months (Figure 5A). Through quantification of the tdTomato reporter gene and qPCR, we confirmed overexpression of miR-223 with AAV uptake mostly in neurons, however we also observe doverlap between tdTomato signal and IBA1 in the peri-plaque area (Figure S8A-B). On behavioral assays, AAV-miR223 treatment did not alter performance in the open field test (Figure S8C). On the Barne’s maze learning days, miR-223 reduced latency to the target hole in App^NL-G-F^ mice, with a trend towards decreasing in WT mice. (Figure 5B, S8D). On the probe test, App^NL-G-F^ mice performed significantly worse than WT mice, however AAV-miR223 partially attenuated this deficit in App^NL-G-F^ mice (Figure S8E). In the contextual fear test, WT mice treated with AAV-miR223 were significantly more active at baseline during training days, however all mice were able to associate the tone and shock, with WT AAV-miR223 mice having a downward shifted learning curve (Figure S8F). On the tone and context test days, performance between App^NL-G-F^ mice with and without miR-223 was not significantly changed (Figure S8F-H). Consistent with the learning trials, WT mice treated with AAV-miR223 spent less time freezing than the WT mice treated with the scrambled oligonucleotide (Figure S8F-H).

**Figure 5:**
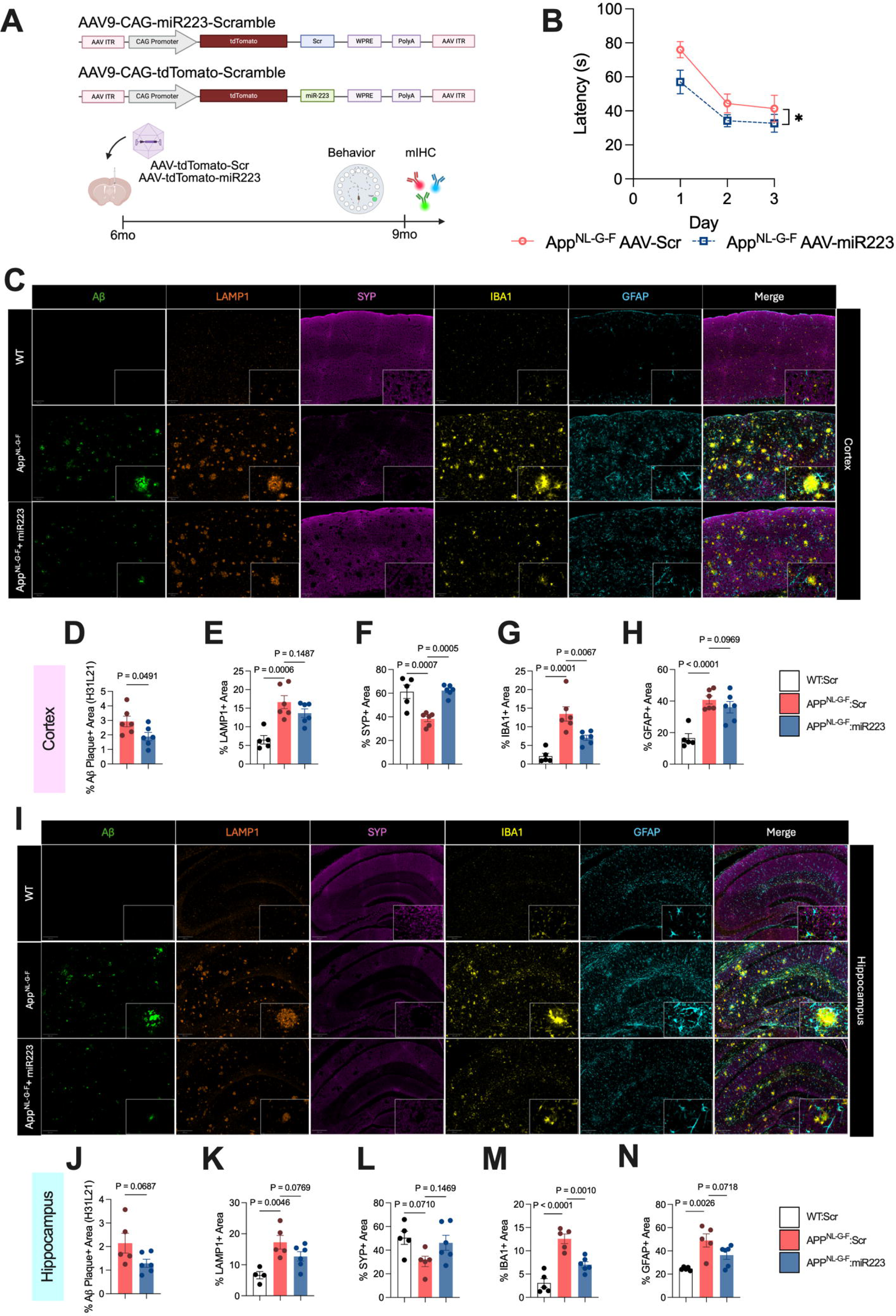
Long-Term AAV-Mediated Overexpression of miR-223 Alleviates Gliosis and Amyloid Pathology. **A** AAV construct map and experimental design for animal studies. **B** Barnes maze latency to hole reveals significantly improved learning in App^NL-G-F^ mice treated with AAV-miR223 relative to scrambled control. **C** Representative IHC images in the cortex for Aβ, LAMP1, SYP, IBA1, and GFAP. **D-H** Quantification in the cortex of Aβ (**D**), LAMP1 (**E**), SYP (**F**), IBA1 (**G)**, and GFAP (**H**) demonstrates reduction in Aβ pathology and restored levels of inflammatory and synaptic markers. **I** Representative IHC images in the hippocampus for Aβ, LAMP1, SYP, IBA1, and GFAP. **J-N** Quantification in the hippocampus of Aβ (**J)**, LAMP1 (**K**), SYP (**L**), IBA1 (**M)**, and GFAP (**N**) demonstrates reduction in Aβ pathology and restored levels of inflammatory and synaptic markers. Data shown as mean +/− SEM. **B** Repeated measures two-way ANOVA **E, J** two-tailed t-test. **E-H, K-N** one-way ANOVA with Holm-Sidak post-hoc test.

We next used IHC to assess AD-related protein levels, including Aβ, IBA1, GFAP, LAMP1, as well as SYP. In the cortex, we found that AAV-miR-223 reduced the levels of Aβ (Figure 5C-D), replicating the oligonucleotide treatment data. Comparing App^NL-G-F^ mice receiving AAV-Scr and AAV-miR223, we observed a trend towards decreased LAMP1+ area (Figure 5E), as well as restored SYP levels (Figure 5F). Measuring markers of gliosis, we found that long-term miR-223 overexpression decreased levels of IBA1 and trended towards a decrease in GFAP (Figure 5G-H). In the hippocampus, there was a similar trend towards decreased Aβ. (Figure 5I-J). LAMP1 and SYP levels likewise had a trend towards reversal of changes in AAV-miR223 treated mice relative to App^NL-G-F^ however this was not statistically significant (Figure 5K-L). Gliosis markers IBA1 was significantly decreased by AAV-miR223 treatment while GFAP had a trend towards decreasing, demonstrating resolution of inflammation (Figure 5M-N). In WT mice treated with AAV-miR223, we found a trend towards increased IBA1, as well as increased LAMP1 staining in the cortex relative to WT AAV-Scr controls (Figure S8I-J), however observed no changes in Synaptophysin or GFAP in either brain region. Notably, increased IBA1+ area in WT AAV-miR223 was not associated with amoeboid phenotype, but rather increased density of processes.

Moreover, in WT mice AAV-miR223 was not associated with increased number of microglia (Figure S8K). Taken together, these data show subtle changes in microglia unique to WT AAV-miR223 mice, possibly relating to their altered behavior in the Barnes maze and CFC tests. Overall, we found that AAV-mediated overexpression of miR-223 recapitulated many of our previously observed phenotypes, supporting its potential as a long-term therapeutic for AD.

## Discussion

Overall, we have shown that miR-223 is protective in AD model mice primarily by promoting microglial-mediated clearance of Aβ, an effect that is accompanied by the preservation of the brain neuronal and synaptic markers and amelioration of behavioral deficits. These actions of miR-223 occurred equally in female and male mice. miR-223 has been mostly studied in the periphery, where it has anti-inflammatory functions in diverse diseases such as cancer, intestinal inflammation, and atherosclerosis^44–48^. In the brain miR-223 is produced primarily by glial cells, however it is often secreted in exosomes or bound to high density lipoprotein (HDL), where it can be taken up by neurons and contribute to signaling in these cells^23–25,49,50^. In models of subarachnoid hemorrhage, miR-223 is transferred from microglia to neurons via exosomes where it contributes to neuroprotection^51^. miR-223 has also been identified in neuron-derived vesicles, and its levels are reportedly decreased in schizophrenia^52^. These features uniquely position miR-223 as a highly pleiotropic miRNA that is protective in multiple cell types. This endogenous broad specificity may be beneficial in the development of therapeutics for complex diseases like AD, where multiple cell-types are dysfunctional in tandem to contribute to disease onset.

Our current study showed that miR-223 consistently and rapidly reduced Aβ pathology in the cortex and hippocampus and requires microglia to perform this function. Our *in vitro* data using human iMGs demonstrated that both miR-223 overexpression and knock down directly influence microglial phagocytosis, indicating that miR-223 is a major regulator of microglial phagocytic capacity. These data are in line with studies in peripheral tissues, such as the liver, adipose, and heart, where miR-223 promotes resident macrophage phagocytosis and anti-inflammatory polarization, ultimately fostering tissue repair^53^. In the CNS, miR-223 enhances clearance of myelin debris in models of spinal cord injury and multiple sclerosis^21,54^. Interestingly, a recent study profiling microRNA expression in 5xFAD mice found that miR-223 was among the microRNAs downregulated in Aβ-laden phagocytic microglia relative to non-phagocytic microglia^55^. Placing these data within the context of our current study demonstrating that the levels of miR-223 are reduced in brain of App^NL-G-F^ mice and that miR-223 is a positive regulator of phagocytosis suggests that phagocytic uptake of Aβ could contribute to a mechanism that leads to the downregulated expression of the miR in the AD model mice. Mechanistically, we found upregulated expression of several proteins involved in microglia phagocytosis, including TREM2, AXL, and CD11c. While the levels of each protein were modestly changed, the consistent upregulation of several phagocytosis-related receptors together likely enhances to the overall plaque clearance capacity by microglia, though this apparent clearance may also involve plaque compaction^56,57^. TREM2 agonism has recently been explored as a potential therapeutic avenue in AD, based on its essential role in mediating the microglial response to restrict Aβ pathology^58^. Interestingly, CD11c was consistently upregulated in both the cortex and hippocampus of miR-223 treated mice, suggesting an important role in mediating the anti-amyloid action of miR-223. CD11c+ microglia are associated with uptake and lysosomal degradation of Aβ in mouse models of amyloidosis^36^. Notably, CD11c+ microglia were also enriched in the vicinity of plaques of patients treated with anti-Aβ immunotherapy, in line with our data showing restricted expression in the peri-plaque area^59^.

Our *in vitro* transcriptomics identified genes associated with the endo-lysosomal network (ELN) as being directly targeted by miR-223 in iMGs. Proper functioning of lysosomes is essential for maintenance of phagocytic function, with Aβ accumulation leading to lysosomal dysfunction^60,61^. Notably, microglia themselves are slow to degrade Aβ and can thus become overwhelmed, creating a feedback loop that impairs further phagocytic uptake^62,63^. Thus, the enrichment of miR-223 target genes within the ELN suggests that miR-223 may enhance phagocytosis by improving downstream lysosomal function. Additionally, improved recycling of phagocytic receptors to the cell membrane rather than degradation in endosomes could provide a plausible mechanism for the observed increase in the protein levels of several receptors, whose mRNA levels are notably not affected by miR-223. To ascertain potential genes involved in ELN function, we focused on genes whose mRNA expression responded reciprocally to the mimic and inhibitor treatment reasoning that miR-223 likely plays an essential role in their regulation. One such direct target gene of miR-223 was *SPPL2A,* a lysosomal-localized protease that has been identified as a GWAS-significant AD risk gene^64–66^. *SPPL2A* is involved in the proteolytic processing of *TMEM106B,* linking miR-223 to two AD risk genes^67^. Another notable direct target of miR-223 was *RUBCN*, which is a negative regulator of early-late endosome transition via its antagonistic action on Rab7 recruitment^68^. It should be noted that due to the relatively small effect size of miRNAs on individual target genes, the overall effect of miR-223 is likely mediated through coordinated actions of several genes within the same pathway, rather than a single gene.

The reduction in Aβ plaque burden evoked by miR-223 was associated with the rescue of a subset of behavioral impairments in App^NL-G-F^ mice, with a concomitant increase in the levels of several synaptic markers in the cortex and hippocampus. This effect was also recapitulated in human iPSC-derived neurons, where synaptic mRNA species such as *DLGAP2*, *SHANK2*, and *GRIA1* were upregulated in mimic treated cells. Interestingly, in the presence of remaining Aβ pathology, miR-223 treated mice were able to maintain SYP levels and had reduced LAMP1+ dystrophic neurite staining, however it is unclear whether this is due to a direct action of miR-223 on neurons, or a consequence of improved microglial handling of plaques leading to neuroprotection. Previous studies using stroke models have shown that miR-223 directly protected neurons during excitotoxic injury^6^. Additionally, miR-223 protected neurons *in vitro* from IL8-induced injury and increased the expression of neurotrophic factors *in vivo*^69^. With respect to AD, miR-223 was able to prevent Aβ-induced apoptosis in a cell model via targeting the PI3K-Akt pathway, establishing that miR-223 can be directly protective against Aβ^24^. Future studies investigating cell-type specific overexpression of miR-223 in neurons and microglia are required to determine the contributions of each cell type to the overall neuroprotective phenotype.

Using an AAV-mediated approach, we found that long-term (3-month) overexpression of miR-223 recapitulated several of the phenotypes previously observed with a short-term treatment, such as decreased Aβ burden, reduced reactive gliosis, and increased synaptic marker expression. Despite long-term exposure to miR-223, the reduction of amyloidosis due to AAV9-mediated overexpression was relatively comparable to short-term LNP-mediated overexpression. This could be due to decreased transgene expression over time coupled with the increase in pathology due to age. Additionally, while AAV9 has been shown to target microglia, transgene expression can vary considerably^70^. Conversely, LNP-based approaches have been shown to robustly target RNAi molecules to microglia *in vivo*^71^. As such, the comparable efficacy between the two approaches may be due to a tradeoff between microglial targeting efficiency and exposure time, with LNP-based approaches potentially achieving higher microglial transfection despite the shorter time frame.

There are several limitations to our study. Firstly, while miR-223 is effective in mediating the clearance of Aβ in the App^NL-G-F^ model, it is unclear whether miR-223 can effectively modulate tau pathology, the major driver of cognitive impairment in AD. Microglia have different responses to Aβ and tau, with the latter leading to an antigen-presenting state in microglia different from the pro-phagocytic state adopted in response to Aβ^72^. While we have shown that miR-223 alters microglial phagocytosis, it is not clear whether it alters tau-associated states. Thus, overexpression of miR-223 in models of tauopathy could further help to evaluate its potential as an AD therapeutic. Additionally, our study only assessed whether miR-223 could alter AD pathology starting at relatively early time points and mild disease (6-7 months). As such, it is unknown whether miR-223 will retain its efficacy in more advanced disease, where microglia begin to show senescence and thus may not be able to effectively respond to miR-223^73^. Lastly, the effects of miR-223 in healthy WT mice is not fully clear. Using oligonucleotide-based overexpression we found no change behaviorally in WT mice treated with miR-223. However, when using AAV-mediated overexpression we found an improvement in latency to find the escape hole in the Barne’s maze accompanied by a paradoxical decrease of freezing in the CFC. WT-AAV-miR223 mice were more active before receiving any shock but still learned to associate the tone with the shock, suggesting that the poor performance on the CFC assay is a consequence of altered baseline activity rather than impaired learning per-se. Notably, decreased freezing was not found in App^NL-G-F^ mice treated with AAV-miR223, suggesting this effect is specific to WT mice. Additionally, we found in WT AAV-miR223 there was an increase in IBA1 staining area, however it is not clear whether the effect on miR-223 on CFC performance is mediated by microglia. Future work understanding the basic biology underlying miR-223 in healthy mice will be essential in determining its safety as a potential therapeutic.

In this study we used a miRNA mimic to treat AD-like phenotypes in a mouse model with the idea that it can recalibrate an entire disease-related dysregulated gene expression network (such as neuroinflammation) as one miRNA coordinately tunes multiple transcripts. Intrathecal delivery of antisense oligonucleotide (ASO) therapeutics, which could be considered as analogous to microRNAs is already successfully used humans, such as nusinersen, for spinal muscular atrophy, and tofersen, for SOD1-associated amyotrophic lateral sclerosis^74^. Most relevant to the current study, encouraging new data on diranersen (a MAPT-targeting, tau-lowering ASO) point to the feasibility of RNA-based therapies for AD as well. Our data support the use of miR-223 as a potential therapeutic for AD and highlight the promise of miRNA-based gene network-targeting therapeutics for neurodegenerative diseases.

## Methods

### Animal Ethics and Care

All animal studies were approved by the Boston University Institutional Animal Care and Use Committee. C57BL6/J (Charles River Laboratories) wild type or App^NL-G-F^ mice (RIKEN) were like sex group housed on a reverse light cycle (11PM-11AM lights on) and had free access to standard chow and water.

### Lipid Nanoparticle Preparation

Oligonucleotides for *in vivo* and *in vitro* studies were purchased from Qiagen (miRCURY LNA miRNA Mimics/Inhibitors). microRNA mimics, inhibitors, and scrambled controls were prepared using the Vivofectamine LNP system. Each animal dose was prepared by mixing 1 µL of 0.5 mM oligonucleotide in sterile nuclease free water with 0.5 µL Vivofectamine reagent and 0.5 µL of 120 mM Sodium Acetate in sterile water. LNPs were prepared by hand by pipetting up and down at least 20 times and were incubated for at least 1 hour prior to injection. The final concentration oligonucleotide in LNP solutions was 0.25 nmol/µL.

### Adeno-Associated Virus Preparation

AAVs using the pAAV-CAG-tdTomato AAV construct (Addgene #59462) were prepared by GeneWiz. Sequences containing +/− 200 nt of the mouse miR-223 hairpin sequence, or a non-targeting control were cloned into the AAC plasmid in the 3’UTR between the EcoRI and HindIII restriction sites. Approximately 1×10^11^ viral particles were administered in 2 µL per mouse.

### Intracerebroventricular Injections

For intracerebroventricular injections, mice were anesthetized using 2% isoflurane in oxygen and were positioned in a stereotaxic instrument. Prior to surgery, mice were injected subcutaneously with 0.05 mg/kg buprenorphine for pain management. A small incision was made on the skull and a small hole in the bone was made using a 27-gauge needle. Mice were injected at the following stereotaxic coordinates relative to bregma: AP −0.6 mm, ML 1.2 mm, DV −2 mm. A total of 2 µL of LNP or AAV solution was injected at a rate of 1 µL/min into the lateral ventricle using a 5 µL Hamilton syringe. The injection apparatus was left in for at least 2 minutes before slowly removing to minimize reflux. Immediately following surgery, mice were injected with 30 mg/kg of ampicillin, 5 mg/kg of meloxicam, and 1 mL of saline to prevent dehydration. Mice were allowed to recover on a heating pad before being returned to their home cage. The day following surgery, mice were given a dose of buprenorphine and meloxicam to aid recovery. Two days post-surgery, mice were given a dose of buprenorphine. Mice were given 5 days to recover following surgery prior to behavioral testing.

### Microglial Depletion with PLX5622

Mice were provided ad libitum access to Open Standard Diet (OSD) control or containing 1200 ppm PLX5622 (D19101002; Research Diets Inc) for 7 days prior to surgery. Following surgery, mice remained on the PLX5622 or control OSD until euthanasia.

### Behavioral Testing

Mice were given 30 minutes prior to starting behavioral testing to acclimate to the room. Mice were temporarily single housed during behavioral testing. Male mice were always tested before female mice. Tracking was performed using the Ethovision XT 14 software. Mice were given 2 rest days between different test batteries.

#### Open Field Test

Mice were placed in a 2700 cm^2^ plexiglass arena for 15 minutes and allowed to freely explore. The center was recognized as the central 1000 cm^2^ region and times spent in the center and periphery were quantified.

#### Barnes Maze

Mice were placed on a circular elevated platform of diameter 92 cm with 20 evenly spaced holes along the periphery in a well-lit room with white noise being played. An escape pod was placed under the same hole for each mouse on each day. Salient visual cues for orientation were placed on the walls near the maze. The test was conducted over the course of 5 days. On day 1, mice were given 2 minutes to acclimate to the test and were gently nudged towards the escape hole if they were unable to enter it. On days 2-4, mice were each given 3 trials of 90 seconds to enter the escape hole and were guided to the hole if unable to enter. Each trial was separated by 20 minutes. Mistakes were counted as non-target hole visits. Measurements were taken until mice were able to first find the hole, and the trial was then run until the mice entered the hole or time ran out. Any non-target visits after the mouse found the target hole were not counted as mistakes. On day 5, the escape hole was covered, and mice were given 30 seconds to find the hole as a test.

#### Contextual Fear Conditioning (CFC)

CFC was conducted over 3 days. On day 1, mice were placed in the conditioning chamber with white walls and a wire floor. After 2 minutes of habituation, mice were played a 20 second 2000Hz tone, with a 0.5 mA current passed through the floor grid for 2 second to deliver a mild foot shock during the final 2 seconds of the tone. Mice were given 100 seconds of rest before the next tone and shock. In total, mice received 5 tone-shock pairs. On day 2, mice were placed in the same chamber with stripped walls and a solid floor. After 2 minutes of habituation, the same 20-second tone was played without a foot shock. A total of 5 tones were played with a 100 second interval between tones to assess memory of the tone. On day 3, mice were placed in the conditioning box with the original white walls and wire floor for 5 minutes with no tone or shock to assess contextual memory. Time spent frozen (i.e. immobile) was measured as the main readout of fearful memory.

### Tissue Extraction

All mice were euthanized using CO_2_ asphyxiation followed by secondary decapitation. The right hemisphere was quickly fixed in periodate-lysine-paraformaldehyde (PLP) fixative (10 mM sodium periodate, 4% paraformaldehyde, 75 mM lysine) for 24 hours at 4 ℃ and transferred to 10% and 20% glycerol in 2% DMSO/0.1M PBS on the first and second days, respectively. The left hemisphere had the hippocampus and frontal cortex micro dissected and snap-frozen on dry ice for downstream assays.

### Cell Culture

#### hiPSC-Derived Neurons

Human iPSCs harboring a PiggyBac Tet-On NGN2 construct (BU2 and BU3 lines, provided by the BU Center for Regenerative Medicine) were maintained on matrigel-coated plates (Corning #356234) using mTeSR Plus media (StemCell Technologies # 100-0276) with daily feeds. iPSC-derived neurons (iNeurons) were differentiated using the protocol described in Ref.^75^. Briefly, iPSC neurons were seeded at ∼1 million cells per well in a 6 well plate coated with matrigel. The initial stages of differentiation were carried out in DMEM/F12 media with N2 supplement (Gibco #17502048), L-glutamine (Gibco #25030081), non-essential amino acids (NEAA, Gibco #11140050) and 2 µg/mL doxycycline (StemCell Technologies #72742), with daily media changes for 3 days. On day 3, cells were dissociated with accutase (StemCell Technologies #07992) and replated on poly-D-lysine (Gibco #A3890401)/matrigel coated plates at a confluency of 100-200k cells/cm^2^, depending on the application. iNeurons were matured in Brainphys Media (StemCell Technologies # 05790) supplemented with B27 (ThermoFisher #17504044), BDNF (PreproTech # 450-02), NT3 (PreproTech # 450-03), and 50 µg/mL laminin (Gibco #23017015). Neuron identity was based on mRNA and protein expression of neuronal marker MAP2.

#### hiPSC-Derived Microglia

Human induced pluripotent stem cells were differentiated into microglia using the STEMdiff™ Hematopoietic, Microglia Differentiation, and Microglia Maturation Kits (STEMCELL Technologies, Catalog #05310, #100-0019 and 100-0020) according to the manufacturer’s instructions. Briefly, iPSCs were passaged as cell aggregates (approximately 100–200 µm diameter) and seeded onto Matrigel-coated plates at a density yielding 16–40 adherent colonies per well after 24 h. Hematopoietic differentiation was induced through a two-stage protocol involving sequential culture in STEMdiff™ Medium A and Medium B. Non-adherent hematopoietic progenitor cells were harvested on day 12 and plated onto Matrigel-coated 6-well plates in STEMdiff™ Microglia Differentiation Medium at a density of 1–2 × 10^5^ cells per well.

Cells were cultured for 24 days with medium supplementation every other day. On day 24, cells were transferred to fresh Matrigel-coated plates and cultured in STEMdiff™ Microglia Maturation Medium for an additional 4–10 days to generate terminally differentiated microglia. Mature microglia were used for downstream functional assays and molecular analyses between days 28 and 34 of differentiation. Cell identity and differentiation efficiency were assessed by staining for IBA1 as recommended by the manufacturer.

#### Cell Transfection with miRNA Mimics and Inhibitors

miRNA locked-nucleic acid mimics (Qiagen #339173) were dissolved to a stock concentration of 10 µM in nuclease free water. miRNA 2’O-methyl antisense inhibitors (IDT) were dissolved to a stock concentration of 10 µM in nuclease free water. Unique scrambled controls for mimic and inhibitor studies *in* vitro were used due to the different chemistries of the oligonucleotide (LNA for mimic, 2’-O-methyl for inhibitors). Transfections were performed using the Lipofectamine RNAiMAX transfection reagent (Invitrogen #13778075) at a final concentration of 30 nM for 48 hours before collection.

#### Intracellular Calcium Flux assay

Intracellular Ca²⁺ dynamics were measured using the Fluo-4 NW Calcium Assay Kit (Thermo Fisher Scientific, # F36206) according to the manufacturer’s instructions with minor modifications. Human iPSC-derived neurons were plated in poly-D-lysine/laminin and martigel coated 96-well plates at a density of 80,000–120,000 cells per well. Neurons were transfected with mimics or inhibitor for 48 hours prior to the assay. For dye loading, a 1× Fluo-4 NW dye solution was prepared in assay buffer (1X HBSS with Ca²⁺/Mg²⁺ + 20 mM HEPES pH 7.4) supplemented with 2.5 mM probenecid to prevent dye extrusion. The cells were incubated with 100 µL dye solution per well for 30 min at 37°C followed by 30 min at room temperature. Fluorescence was recorded using a Varioskan LUX with excitation/emission settings of 494/516 nm. Baseline fluorescence was acquired for ∼2 min, followed by stimulation with glutamate (100 µM). Changes in fluorescence were monitored for an additional 5–30 min.

Fluorescence data were analyzed as normalized changes in signal (ΔF/F₀), where F₀ represents the mean baseline fluorescence intensity measured prior to stimulation and ΔF = F − F₀. Traces were background-corrected and expressed as ΔF/F₀ to account for variability in dye loading and cell number. Each condition was measured in six technical replicates (n = 6 wells per condition). Data are presented as mean ± SEM. Statistical comparisons between groups were performed using unpaired two-tailed Student’s t-test.

#### Phagocytosis Assay

Microglial phagocytic activity was measured using the Vybrant™ Phagocytosis Assay Kit (Invitrogen, Catalog # V-6694) according to the manufacturer’s protocol. Briefly, cells were seeded in black 96-well plates at 40,000 cells per well and allowed to adhere. Fluorescein-labeled *E. coli* BioParticles were added to each well and incubated for 2 h at 37°C. Extracellular fluorescence was quenched with trypan blue, and internalized particle fluorescence was measured using a fluorescence plate reader at approximately 480 nm excitation and 520 nm emission. Phagocytosis was quantified after background subtraction and normalized to scrambled microRNA treated control wells.

### Immunofluorescence

Free-floating 40 µm mouse brain coronal sections comprising the intermediate hippocampus were cut frozen using a microtome. Sections were blocked using 10% donkey serum and 0.3% Triton X-100 in PBS, followed by quenching with 3% water/50% methanol for 15 minutes. Sections were incubated in primary antibody (IBA1 1:1000 Wako #019-19741, CD68 1:200 CST #17846, CD11c 1:200 CST #97585) in 1% BSA/0.3% Triton X-100 overnight at 4 ℃. Sections were washed 3 times in PBS and incubated with Alexafluor 594 Donkey anti-Rabbit (Invitrogen #A-21207 at 1:2000 in 10% Donkey serum/0.3% Triton X-100 for 3 hours at room temperature. Sections were washed 3 times in PBS. If staining with DAPI, 1:1000 DAPI was added to the first wash. To stain with AmyloGlo (Biosensis #TR-300-AG), sections were incubated in 50% ethanol in distilled water for 5 minutes, followed by washing in distilled water for 2 minutes. Sections were incubated in a 1:100 dilution of AmyloGlo in distilled water for 10 minutes, followed by soaking in PBS for 5 minutes. For Thioflavin S staining, sections were incubated in 0.01% ThioS in 50% ethanol in PBS for 2 minutes, followed by incubation in 40% ethanol in PBS for 10 minutes. Sections were washed 3 times for 5 minutes in PBS before additional staining. Sections were mounted and coverslipped using Vectashield and sealed using nail polish.

Images were taken on a Zeiss Axioscan slide scanner with a 20x objective using a Z-stack thickness of 1 µm. Cortical and hippocampal regions of interest (ROIs) were selected, and the images underwent maximum intensity projection prior to analysis. Percent area for stains was quantified in QuPath. AmyloGlo stained images underwent background subtraction with a radius of 50 pixels in ImageJ to improve image segmentation, followed by manual thresholding. No additional processing was performed, and all images were processed in the same way. Amyloid plaques were identified and quantified using the analyze particles function in ImageJ. Microglial morphology quantification was performed using the AutoMorFi pipeline for ImageJ^35^. Sliding paraboloid background subtraction with a radius of 200 pixels was uniformly applied to all images to improve image segmentation and detection of fine microglia processes. No other preprocessing was performed. Images were manually thresholded and AutoMorFi was run with a soma size range of 30-2000 and cell size range of 200-5000. Whole sections (1-2 per animal) were analyzed. Researchers were blinded to the treatment group during all analysis steps. Sholl analysis was performed in the SNT ImageJ plugin. Individual microglia were selected in 45 µm x 45 µm analysis windows and the center of the microglial soma was selected as the starting point. One µm step sizes were used for each concentric circle. Representative images underwent minimal processing except for increasing the brightness, which was done uniformly across the whole image and applied to all images.

#### Multiplex Immunohistochemistry

Sections were embedded in paraffin wax and were cut at 5 µm with RM2255 Microtome (Leica, Wetzlar, Germany) for multiplex fluorescent immunohistochemistry (mfIHC). mfIHC was performed on a Ventana Discovery Ultra autostainer (Roche Diagnostics, Indianapolis, IN, USA). Pretreatment was performed with Benchmark Ultra CC1, a Tris-based antigen retrieval buffer, at 95°C for 32 minutes. The secondary antibody for all targets was Vector ImmPress Goat Anti-Rabbit IgG (MP-7451, Vector Laboratories, Newark, CA, USA), predilute solutions incubated for 20 min at 37°C following a protein blocking step with Akoya Opal Diluent/Block (ARD1001EA, Akoya Biosciences, Marlborough, MA, USA). Primary and secondary antibody complexes were developed with Opal fluorophores. A nuclear counterstain was performed with Akoya Spectral DAPI (FP1490, Akoya Biosciences) and slides were mounted with Prolong Gold Antifade Mountant (P36930, Invitrogen, Waltham, MA, USA). The antibodies used were SAP102 (1:100, RT 8h, Opal 690, Invitrogen #PA5-29116), Beta Amyloid (1:100, 37°C 40 min, Opal 520, Invitrogen #700254), ChAT (1:1000, 37°C 40 min, Opal 570, CST #27269), MAP2 (1:1000, RT 1h, Opal 480, Abcam #ab32454), LAMP1 (1:250, 37°C 40 min, Opal 620, CST #46843), SYP (1:100, 37°C 40 min, Opal 780, CST #36406). Whole slide-images were acquired with a PhenoImager HT^TM^ Automated Quantitative Pathology Imaging System and spectrally unmixed on the instrument using InForm (v3.0) (Akoya). Images were analyzed in QuPath.

### Bulk RNA Sequencing

Total RNA was extracted from tissues or cells using the Zymo Quick RNA Miniprep Kit according to manufacturer’s instructions (Zymo #D3024). RNA-sequencing was conducted by SigniosBio. For RNA-sequencing, library preparation was conducted on 200 ng of total RNA using the KAPA mRNA HyperPrep Kit, followed by sequencing on a NovaSeq X plus with a read depth of 20M 150 base pair paired-end reads per sample. FASTQ files were trimmed using Trimmomatic^76^ and reads were aligned and quantified to the mm10 (mice) or hg38 (iPSC-derived neurons) reference genome using Salmon^77^. Differential expression was conducted in R using DESeq2^78^. Gene set enrichment analysis (GSEA) was done using clusterProfiler^79^, and over-representation analysis was done using gProfiler.

### Quantitative PCR (qPCR)

To perform miRNA qPCR, miRNA concentration was quantified using the Qubit miRNA Assay Kit (Q32880) and 40 ng of miRNA was reverse transcribed using the miRCURY LNA RT KIT (339340). cDNA was diluted 1:60 in nuclease free water and qPCR was performed using the miRCURY LNA SYBR PCR kit (339346) using 3 µl of cDNA. RT-PCR was performed on a QuantStudio 12K Flex system. The delta-delta CT method was used for normalization with RNU5G as the housekeeping gene

### Western Blotting

Protein from brain tissue was sonicated in 400 µL of NP40 lysis buffer with 1x protease inhibitor cocktail (Thermo Scientific #78440) and was stored at −80 °C until used. Protein levels were quantified using the BCA assay method and were diluted to 2.5 µg/µL in lysis buffer with 4x loading buffer and 2.5% beta-mercaptoethanol and boiled at 70°C for 10 minutes. Thirty µg of protein was loaded in a 4%-12% polyacrylamide gel (Thermo Fisher #WG1403BOX) and was run at 110V for 10 minutes, followed by 150V for 60 minutes in MOPS or MES buffer. Gels were transferred to PVDF membranes using an iBlot3 dry transfer system and were blocked in Superblock for 1 hour. Membranes were incubated in primary antibody (Axl 1:500 CST #18554, Mer 1:1000 CST #38102, Actin 1:5000 Sigma #A5316, Asc 1:1000 CST #67824, Cd11c 1:1000 CST #97585, Nlrp3 1:500 CST #15101, Trem2 1:1000 CST #55739) diluted in 5% milk in TBST overnight at 4°C Membranes were washed 5 times with TBST for 5 minutes each before incubation with secondary antibody in 5% milk in TBST for 1 hour at room temperature. Blots were then washed 5 times with TBST for 5 minutes each before imaging. Signal was developed using the SuperSignal™ West Pico PLUS Chemiluminescent Substrate (Invitrogen #34580). Band intensity was quantified in ImageJ and was normalized to beta actin.

### Statistical Tests

All statistical tests were used as described in the figure legends. Outliers were determined via the Grubbs test based on an alpha of 0.05.

## Authors’ Contributions

Concept and design: AK, NG, JKB, AF, ID, TJM,

Acquisition of data: AK, NG, UC, WB, CP, OJ, JL, TSG, JC, AO, CL, NC, MK, JT, AMF, TLT, TJM

Data analyses: AK, NG, UC, AMF, JKB, TJM

Interpretation of data: AK, NG, UC, AF, ID, JKB, TJM

Drafting of the manuscript: AK, JKB, TM

Reviewing and editing: AK, NG, UC, WB, CP, OJ, JL, TSG, JC, AO, CL, NC, MK, JT, AMF, TLT, AF, ID, JKB, TJM

## Acknowledgements

The authors would like to thank Cindy X. Zhang (University of Toronto) for the helpful discussions and suggestions.

---

**Figure S1:**
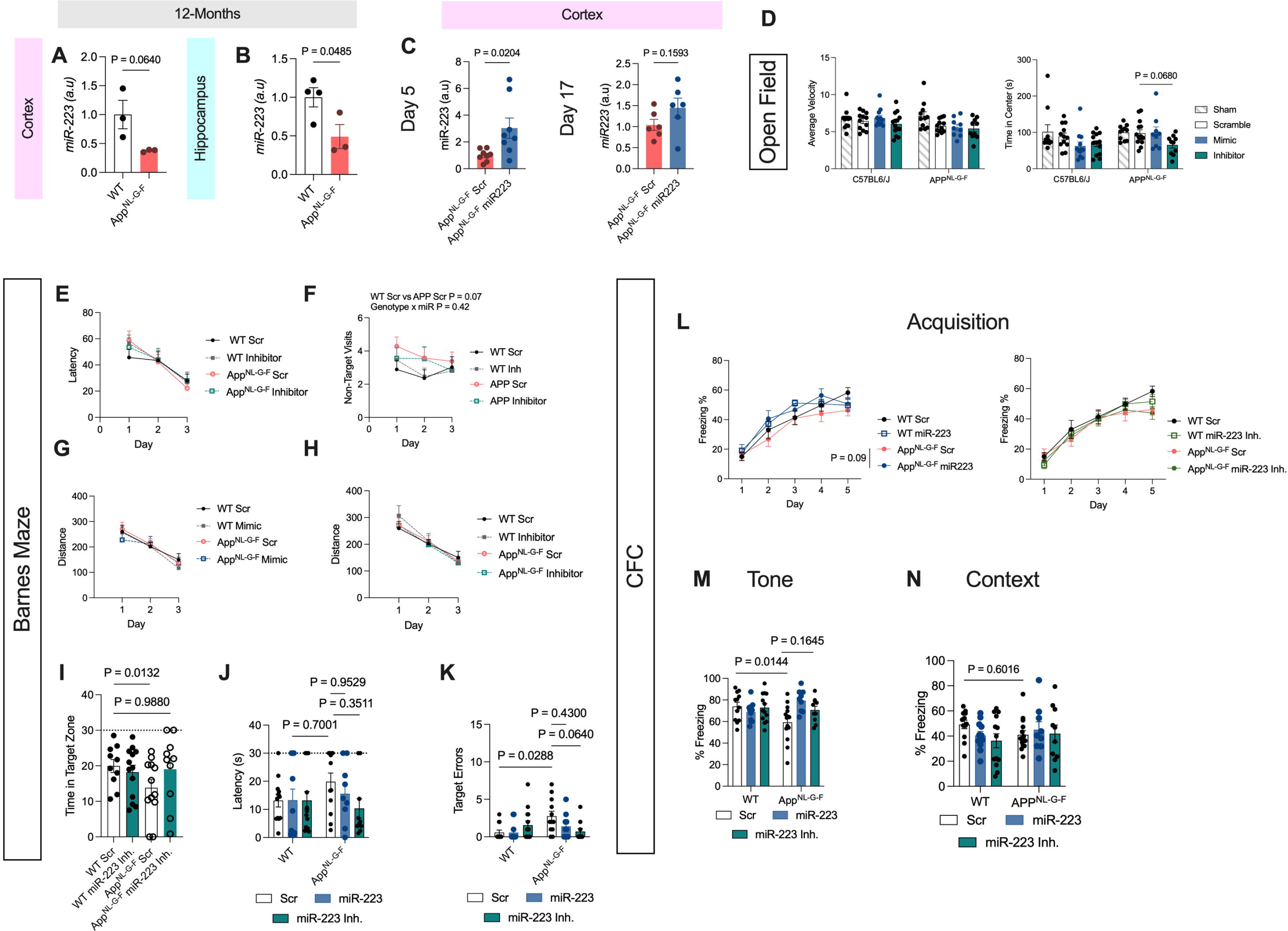
Behavioral Effects of miR-223 Modulation in Mice (Related to Figure 1) **A-B** Validation of miR-223 downregulation in 12-month-old App^NL-G-F^ mice. **C** qPCR of cortical tissue validating miR-223 overexpression at 5 days post-treatment with a trend towards overexpression seen by day 17. **D** Open field test time in center and average velocity demonstrates no change in locomotor activity relative to no surgery (sham) animals, as well as a trend towards increased anxiety in AppNL-G-F mice treated with miR-223 inhibitor. **E-F** Barnes maze training day latency (**E**) and mistakes (**F**) in mice treated with miR-223 inhibitor. **G-H** Path length to the escape hole in mice treated with miR-223 mimic or inhibitor. **I-K** Time in target quadrant (**I**) in mice treated with miR-223 inhibitor on the 1-day probe test and latency to hole in all groups (**J**) and mistakes made (**K**) on the 1-day probe test. **L** Acquisition curves during foot shocks during day 1 of the contextual fear conditioning (CFC) paradigm (left = miR-223 mimic, right = miR-223 inhibitor). **M** miR-223 inhibitor does not change freezing behavior during the tone test relative to scrambled control. **N** miR-223 overall does not change freezing during the context text relative to scrambled control.

**Figure S2:**
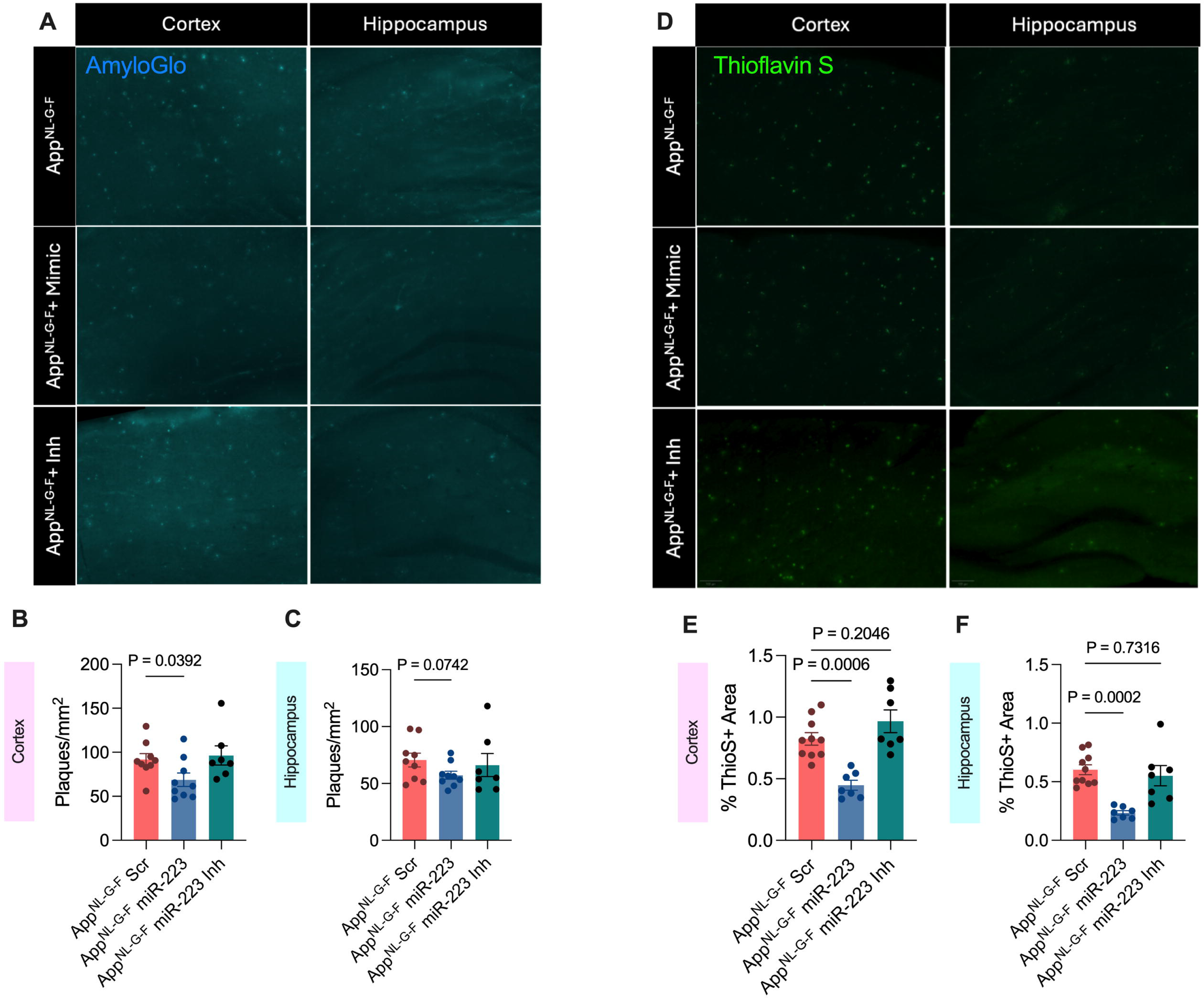
Role of miR-223 on Amyloidosis in App^NL-G-F^ Mice (Related to Figure 1) **A** Representative AmyloGlo staining images of experimental groups in the cortex and hippocampus. **B-C** Quantification of the number of Aβ plaques in the cortex and hippocampus shows significant reduction in mimic treated mice, with no change in inhibitor treated mice. **D** Representative ThioS staining images of experimental groups in the cortex and hippocampus. **E-F** ThioS+ staining in the cortex and hippocampus shows no change with miR-223 inhibitor treatment.

**Figure S3:**
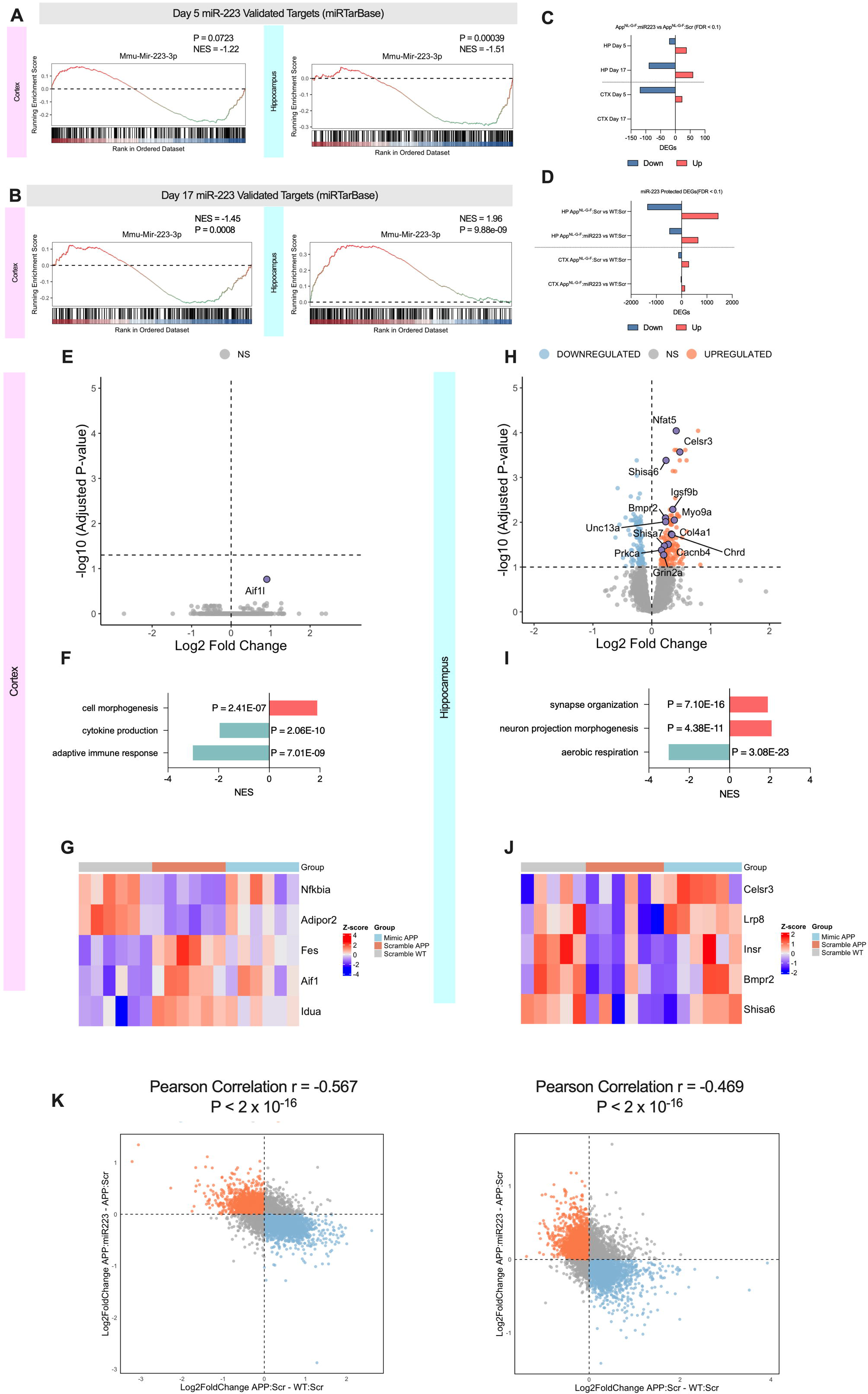
Transcriptomic Characterization of miR-223 (Related to Figure 1) **A** GSEA enrichment of miR-223 validated target genes changed by miR-223 at day 5 in the cortex and hippocampus. **B** GSEA enrichment of miR-223 validated target genes changed by miR-223 treatment at day 17 in the cortex and hippocampus. **C-D** Number of up and downregulated DEGs in the noted treatment groups. **E** Volcano plot of DEGs between miR-223 versus scrambled control treated App^NL-G-F^ mice at 17 days post-treatment in the cortex. **F** Pathway enrichment in the cortex of day 17 miR-223 treated mice reveals downregulation of immune response. **G** Heatmap of miR-223 protected genes in the cortex of day 17 miR-223 treated mice. **H** Volcano plot of DEGs between miR-223 versus scrambled control treated App^NL-G-F^ mice at 17 days post-treatment in the hippocampus. **I** Pathway enrichment in the hippocampus of day 17 miR-223 treated mice reveals upregulation for synapse organization. **J** Heatmap of miR-223 protected genes in the hippocampus of day 17 miR-223 treated mice. **K** Correlation of Log2FoldChange values between miR-223 vs scrambled control in App^NL-G-F^ mice and App^NL-G-F^ vs WT mice in the cortex and hippocampus. The top left quadrant (red) represents genes downregulated in App^NL-G-F^ vs WT mice and upregulated in miR-223 vs scrambled control in App^NL-G-F^ mice. The bottom left quadrant (blue) represents genes upregulated in App^NL-G-F^ vs WT mice and downregulated in miR-223 vs scrambled control in App^NL-G-F^ mice.

**Figure S4:**
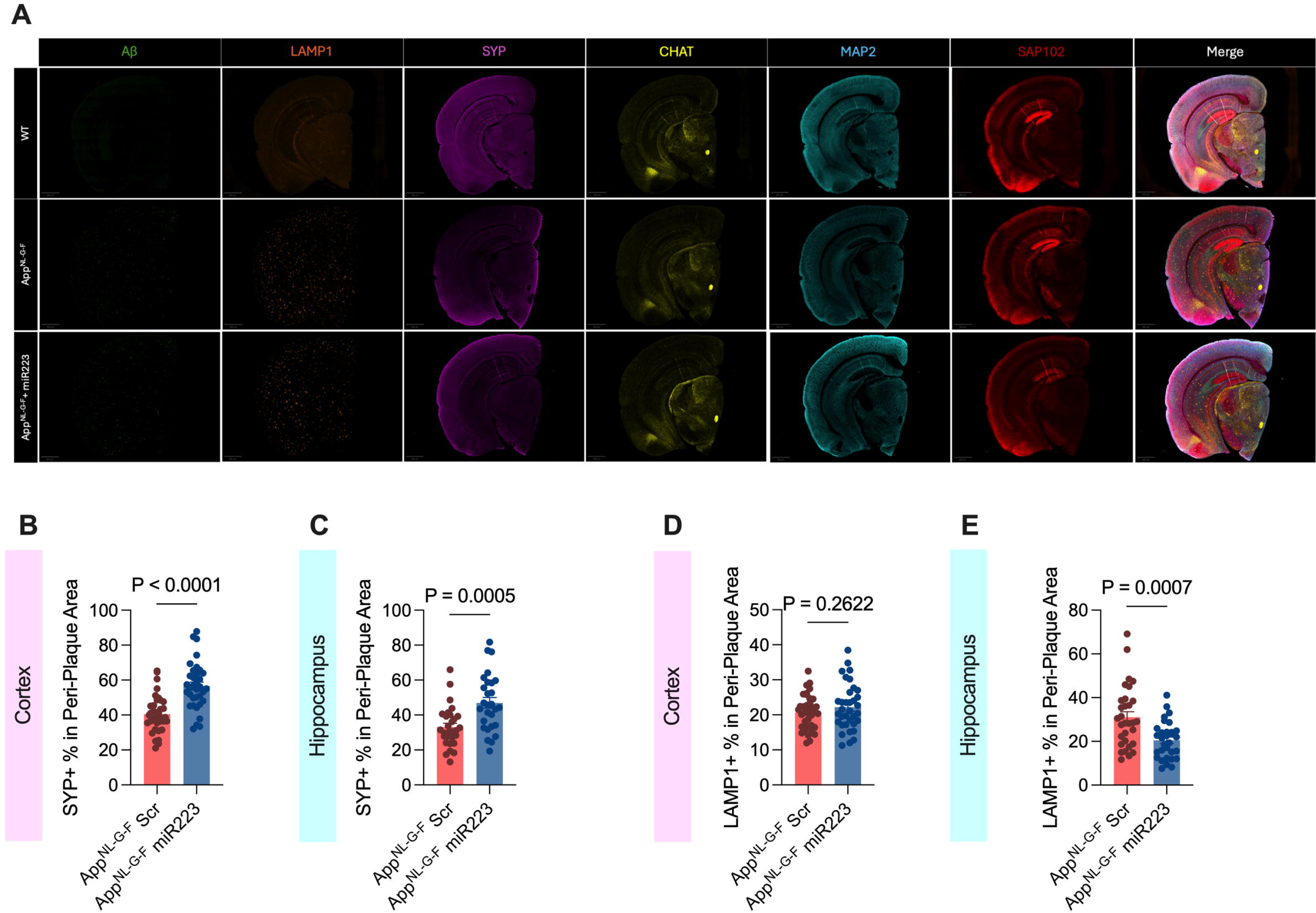
Multiplex Immunohistochemical Staining of miR-223 Treated Mice (Related to Figure 2) **A** Representative whole hemi-brain images of multiplex IHC-staining in wild type scrambled control (WT), App^NL-G-F^ scrambled control and App^NL-G-F^ miR-223 treated mice. **B-C** Distribution of peri-plaque SYP+ area between scrambled control and miR-223 treated mice (n = 30 plaques from n = 6 mice) in the cortex and hippocampus. Each dot represents one plaque. **D-E** Distribution of peri-plaque LAMP1+ area between scrambled control and miR-223 treated mice (n = 30 plaques from n = 6 mice) in the cortex and hippocampus. Each dot represents one plaque.

**Figure S5:**
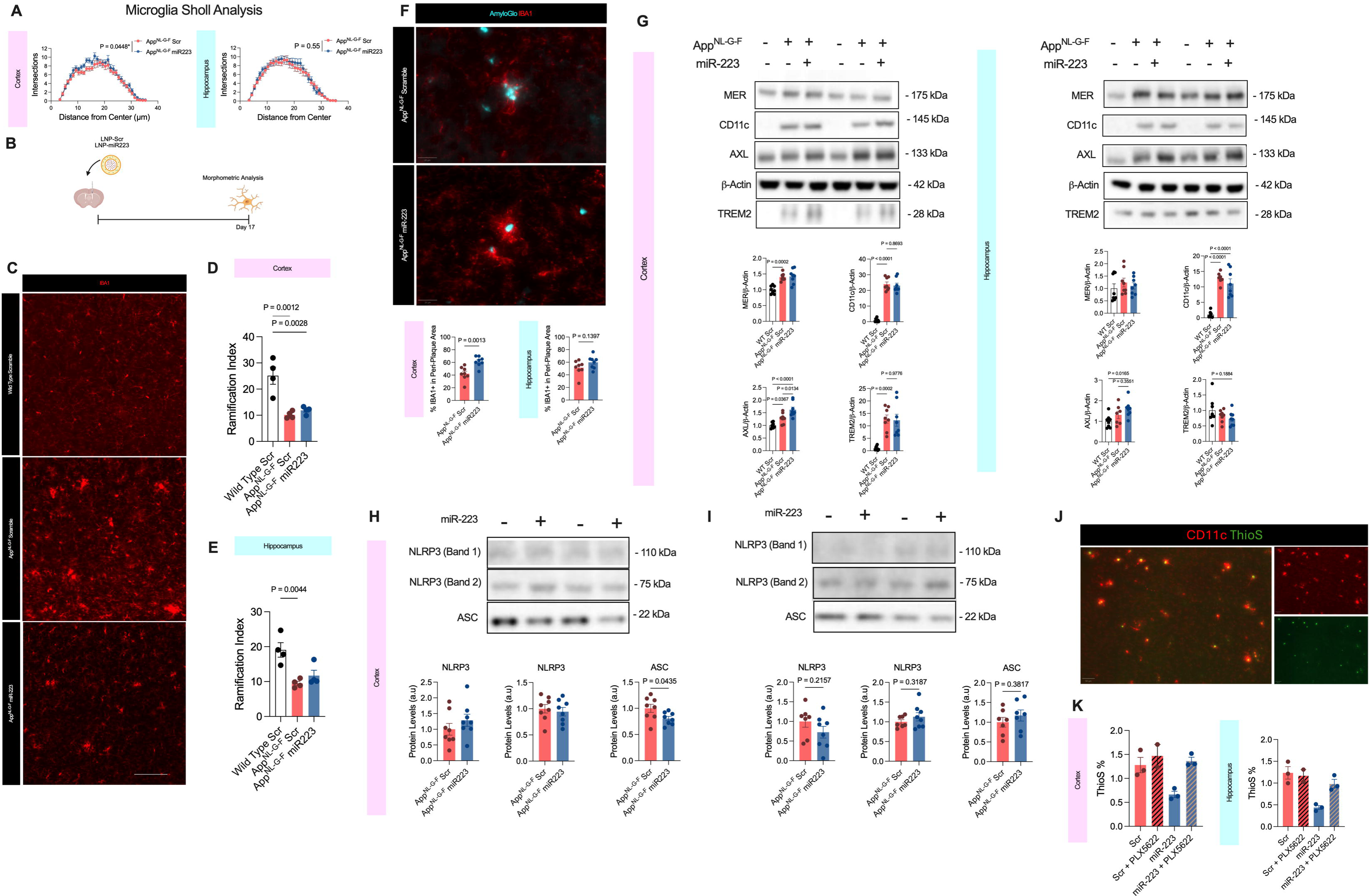
Characterization of Microgliosis in miR-223 Treated Mice (Related to Figure 3) **A** Assessment of microglial ramification via Sholl analysis reveals increased complexity in the cortex of miR-223 treated mice. **B** Treatment time course for assessing microglial changes in miR-223 treated mice. **C** Representative IBA1 staining of day 17 post-treatment WT scrambled control, App^NL-G-F^ scrambled control and App^NL-G-F^ miR-223. **D-E** Ramification index is decreased between WT and App^NL-G-F^ mice and is not changed by miR-223 treatment at day 17 post treatment. **F** Representative images and quantification of peri-plaque IBA1+ area in the cortex and hippocampus at day 17 demonstrates increased clustering around plaques in the cortex. **G** Western blot quantification of microglia receptors involved in Aβ phagocytosis in the cortex and hippocampus of day 17 post-treatment mice, including AXL, MER, CD11c, and TREM2. **H-I** Western blot quantification of NLRP3 and ASC in the cortex and hippocampus of day 5 treated mice. **J** Representative immunofluorescence staining demonstrating restricted expression of CD11c surrounding amyloid plaques. **K** ThioS quantification of amyloid plaques in the cortex and hippocampus shows PLX5622 treatment itself does not alter Aβ pathology after 12 days of treatment.

**Figure S6:**
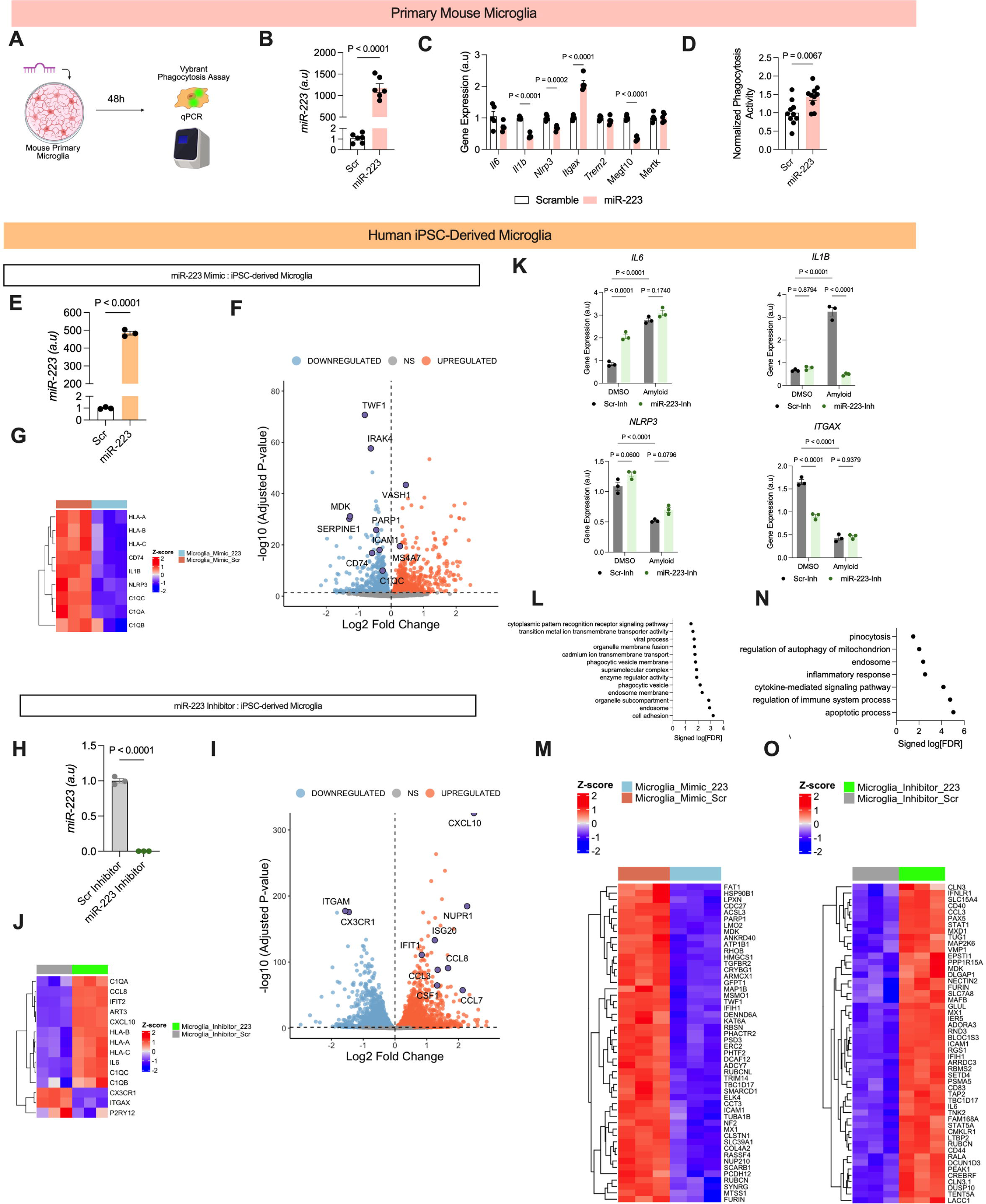
Characterization of miR-223 in iPSC-Derived Microglia (Related to Figure 4) **A** Experimental design for primary microglia studies. **B** Validation of miR-223 overexpression in mimic treated primary microglia via qPCR. **C** Gene expression changes in primary microglia are associated with inflammation and microglial function. **D** Normalized phagocytic activity measured via Vybrant assay in primary microglia treated with miR-223 or scrambled control. **E** Validation of miR-223 overexpression in mimic treated iMGs prior to RNA-seq. **F** Volcano plot of significant DEGs changed with miR-223 overexpression in iMGs. **G** Heatmap of microglial and immune genes downregulated by miR-223 treatment measured via RNA-seq. **H** Validation of miR-223 knockdown in inhibitor treated iMGs prior to RNA-seq. **I** Volcano plot of significant DEGs changed with miR-223 knockdown in iMGs. **J** Heatmap of microglial and immune genes upregulated by miR-223 knockdown measured via RNA-seq. **K** Gene expression for microglial genes *IL6, IL1B, NLRP3*, and *ITGAX* with miR-223 inhibitor and Aβo. **L** Pathway enrichment of direct targets downregulated by miR-223 mimic treatment reveals enrichment of endosome-associated genes. **M** Heatmap of top direct targets downregulated by miR-223 mimic treatment. **N** Pathway enrichment of direct targets upregulated by miR-223 inhibitor treatment reveals enrichment of immune, cytokine, and endosome-associated genes. **O** Heatmap of top direct targets upregulated by miR-223 inhibitor treatment.

**Figure S7:**
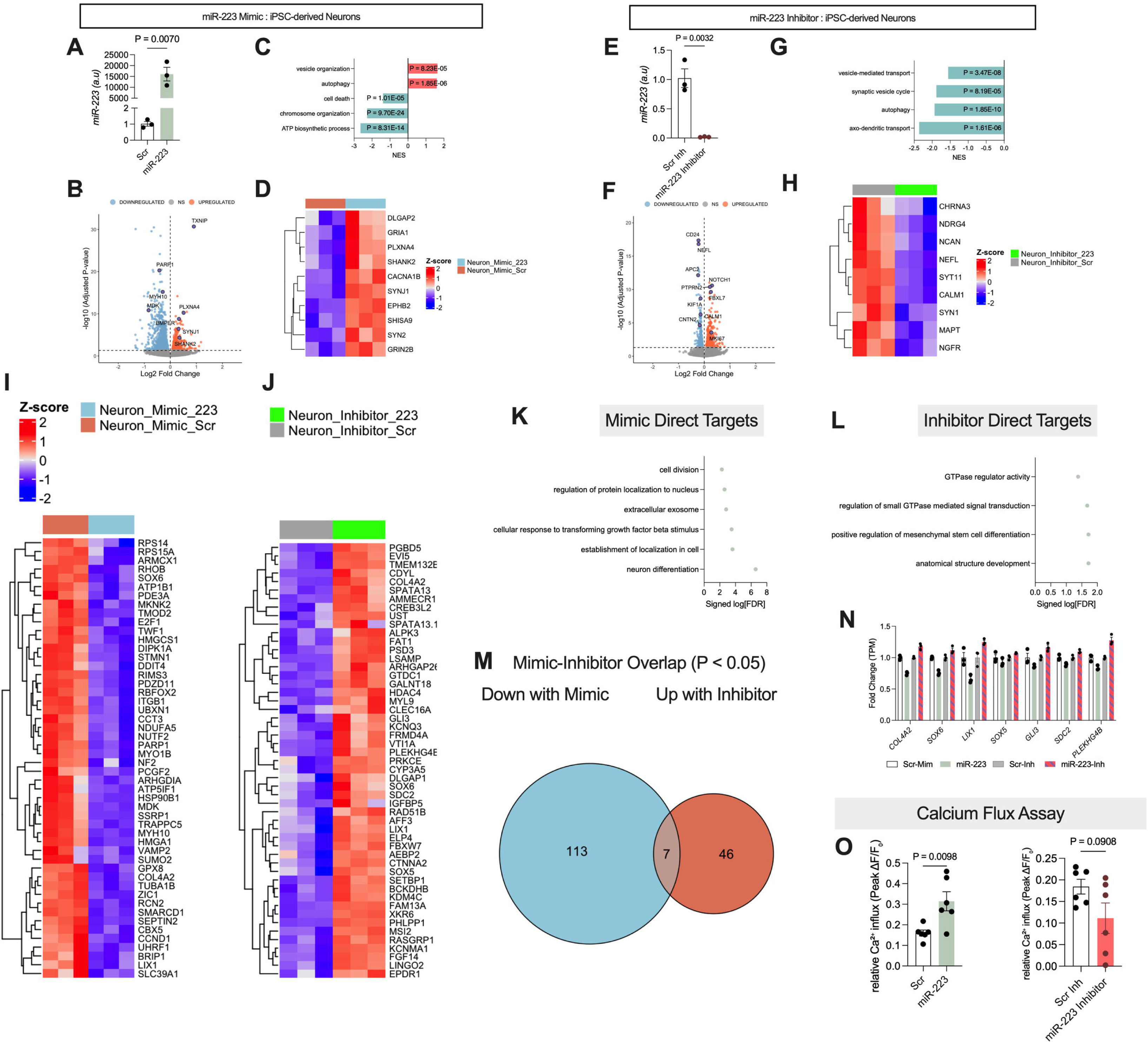
Characterization of miR-223 in iPSC-derived Neurons. **A** Validation of miR-223 overexpression in mimic-treated iNeurons by qPCR. **B** Volcano plot of DEGs changed with miR-223 mimic treatment. **C** Pathway enrichment of top upregulated and downregulated neuronal DEGs upon treatment by miR-223 mimic. **D** Heatmap of synaptic signaling genes and their change with miR-223 treatment. **E** Validation of miR-223 knock-down in inhibitor treated iNeurons by qPCR. **F** Volcano plot of DEGs changed with miR-223 mimic treatment. **G** Pathway enrichment of top upregulated and downregulated neuronal DEGs upon treatment by miR-223 inhibitor. **H** Heatmap of synaptic signaling genes and their change with miR-223 treatment. **I** Heatmap of top miR-223 target genes that are downregulated by mimic treatment in iNeurons. **J** Heatmap of top miR-223 target genes that are upregulated by inhibitor treatment in iNeurons. **K** Pathway enrichment of significantly downregulated target genes with mimic treatment. **L** Pathway enrichment of significantly upregulated target genes with inhibitor treatment. **M** Overlap between direct targets decreased with mimic treatment and upregulated with inhibitor treatment. **N** Fold change RNA-seq values of 7 overlapping genes identified in **M**. **O** Peak calcium flux miR-223 mimic and inhibitor treated iNeurons upon glutamate stimulation.

**Figure S8:**
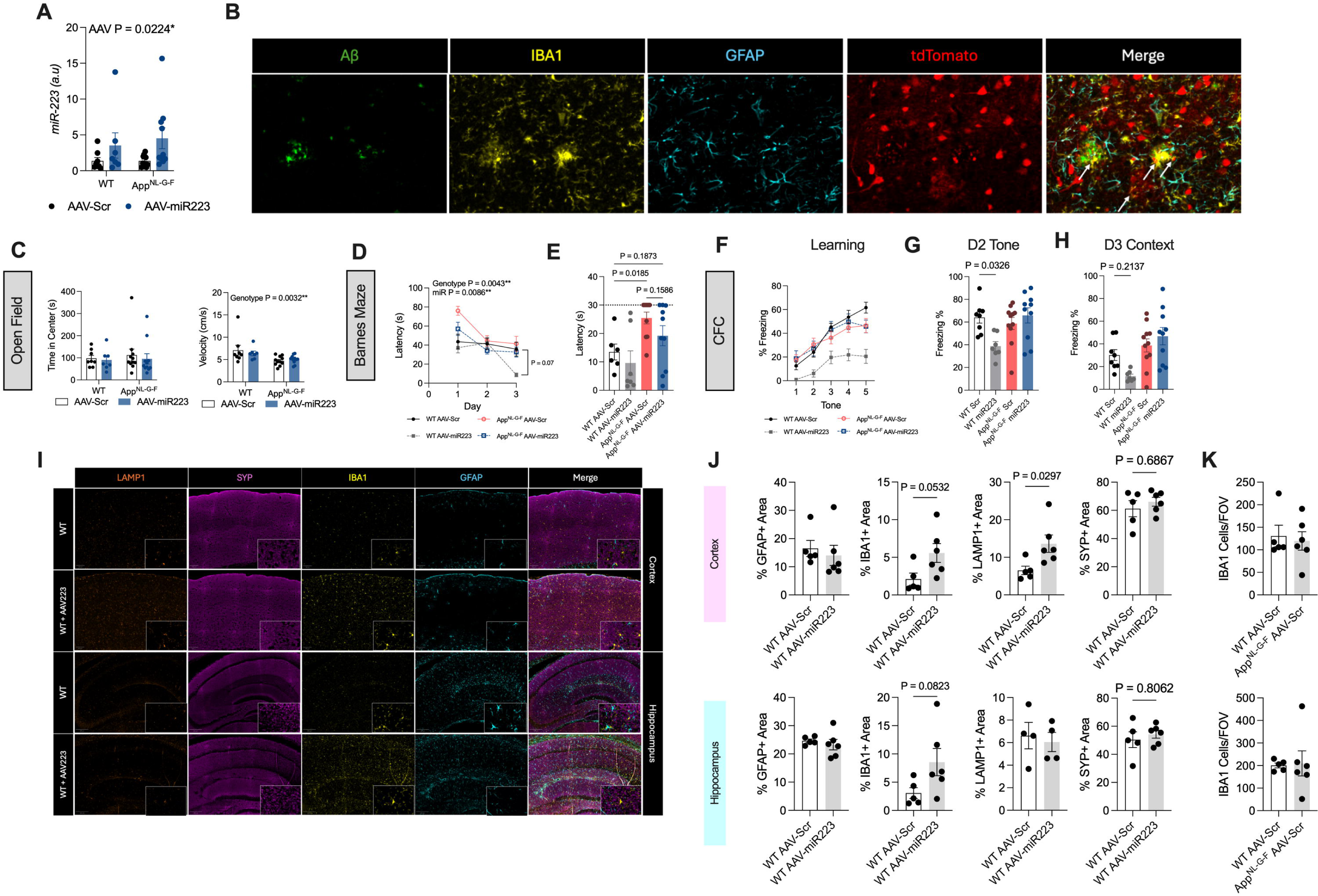
Validation and Behavioral Testing of AAV Mice. **A** qPCR validation of miR-223 levels in AAV mice. **B** Assessment of overlap between tdTomato and glial stains. White arrows represent tdTomato+ microglia. **C** Time in center and average velocity in the open field tests associated with AAV treatment groups. **D-E** Barne’s maze latency to escape hole over trial (**D**) and probe (**E**) days. **F-G** Contextual fear conditioning test across learning, tone and context tests. **I-J** Representative images and quantification of the effect of miR-223 on glial and synaptic markers in WT mice. **K** Microglia numbers per 10 mm^2^ FOV.

